# Tunable phenotypic variability through an autoregulatory alternative sigma factor circuit

**DOI:** 10.1101/2020.06.18.158790

**Authors:** Christian P. Schwall, Torkel Loman, Bruno M.C. Martins, Sandra Cortijo, Casandra Villava, Vassili Kusmartsev, Toby Livesey, Teresa Saez, James C. W. Locke

## Abstract

Genetically identical individuals in bacterial populations can display significant phenotypic variability. This variability can be functional, for example by allowing a fraction of stress prepared cells to survive an otherwise lethal stress. The optimal fraction of stress prepared cells depends on environmental conditions. However, how bacterial populations modulate their level of phenotypic variability remains unclear. Here we show that the alternative sigma factor σ^V^ circuit in *B. subtilis* generates functional phenotypic variability that can be tuned by stress level, environmental history, and genetic perturbations. Using single-cell time-lapse microscopy and microfluidics, we find the fraction of cells that immediately activate σ^V^ under lysozyme stress depends on stress level and on a memory of previous stress. Iteration between model and experiment reveals that this tunability can be explained by the autoregulatory feedback structure of the *sigV* operon. As predicted by the model, genetic perturbations to the operon also modulate the response variability. The conserved sigma-anti-sigma autoregulation motif is thus a simple mechanism for bacterial populations to modulate their heterogeneity based on their environment.

## INTRODUCTION

Cells live in a changeable environment and experience a wide range of environmental stresses. Bacterial populations have evolved strategies to survive these stresses. One strategy is for all cells to immediately respond to stress with the activation of the relevant stress response pathway (Hilker et al. 2016). Alternatively, a bacterial population can generate a broad range of cellular states, which allows it to hedge its bets against the changeable environment (Veening et al. 2008). Noise in gene expression has been proposed as a mechanism for generating phenotypic variability in genetically identical cells (Raj and van Oudenaarden 2008; Martins and Locke 2015). However, how the bacterial population regulates individual cell decisions in order to modulate the fraction of cells that enter an alternative transcriptional state remains unclear (Figure 1A).

**Figure 1.**
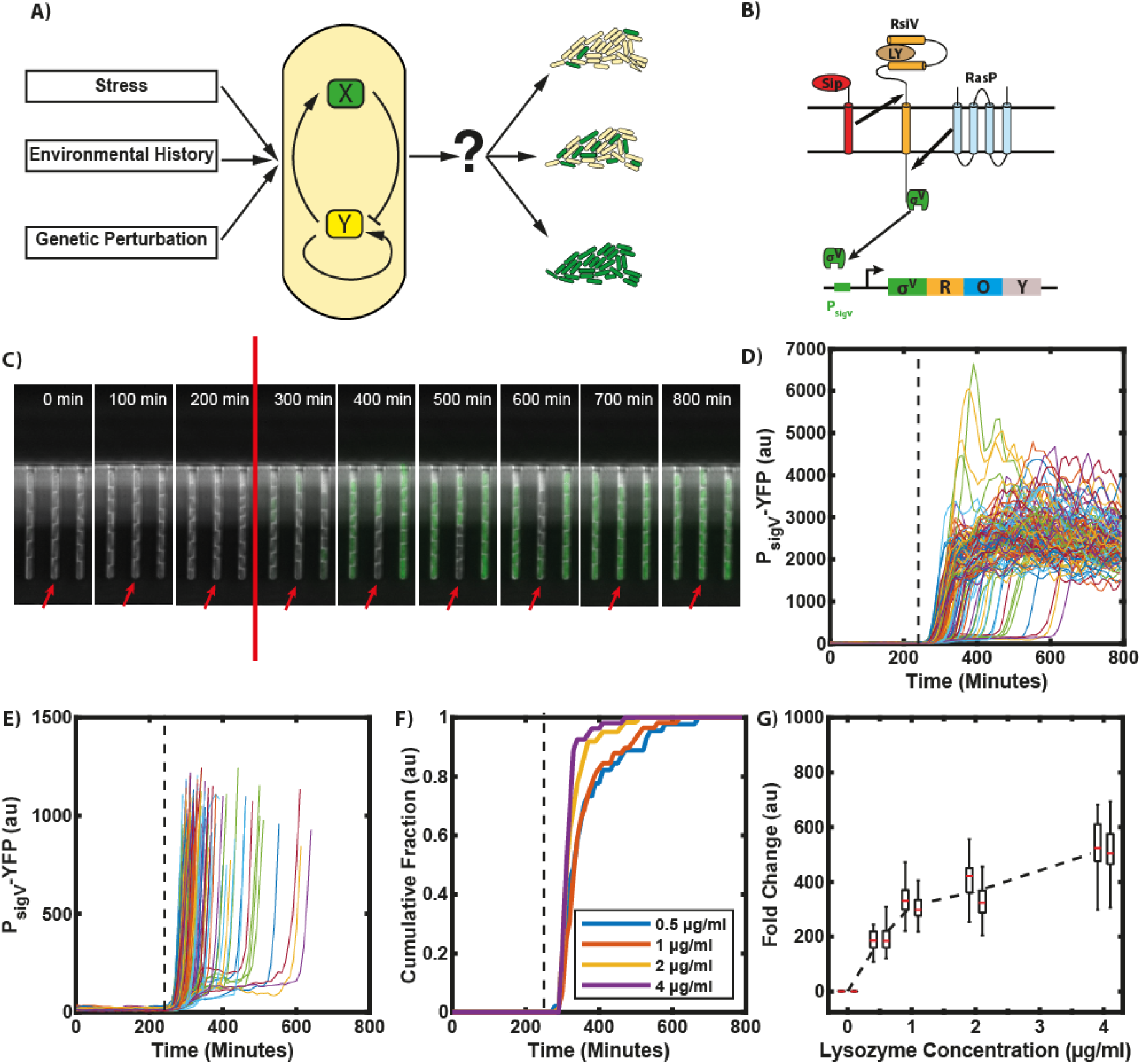
σ^V^ is activated heterogeneously in response to lysozyme stress. (A) It is unclear how genetic circuits (centre) tune phenotypic variability in a bacterial population in response to genetic perturbation, environmental stress and history (left) to modulate the fraction of cells that activate a given pathway (right). (B) Schematic of the σ^V^ circuit. In its inactive form σ^V^ is bound to its anti-sigma factor RsiV. If there is lysozyme present in the environment, RsiV binds to lysozyme and undergoes a conformational change. Only then the proteases SipS and RasP can cleave RsiV to release σ^V^. Once σ^V^ is released from RsiV it can bind to RNA polymerase to redirect transcription to the σ^V^ regulon. σ^V^ can initiate transcription of its own operon which includes *sigV, rsiV, oatA* (one of the main lysozyme resistance genes) and *yrhK* (unknown function). (C) Time-lapse microscopy of cells containing a P_sigV_-YFP promoter reporter reveals heterogeneous activation of σ^V^ in response to lysozyme stress. The stress (red line) was added between the 200 mins and 300 min timepoints (at 240 min). The red arrow highlights a cell with a delayed activation of σ^V^. (D) Time traces of mean YFP fluorescence per cell. Each trace represents a single cell lineage’s response to 1 µg/ml lysozyme. (109 traces from two independent experiments). Lysozyme added at the black dashed line. (E) Figure D replotted to allow examination of variable activation times. (F) The observed heterogeneity is reduced with increasing stress levels. Each line represents the cumulative fraction of cells (N=∼50) with P_sigV_-YFP values that are higher than the half maximum of their final values (representing cells that have activated). (G) The fold change in mean YFP fluorescence increases with increasing stress levels, shown for two biological repeats (adjacent boxplots). Each day’s fold change distribution consisted of >40 manually corrected single cell traces. Error bars correspond to the mean + SD of each repeat.

The σ^V^ mediated lysozyme stress response pathway in *B. subtilis* is an ideal model system to examine how bacterial populations can tune their phenotypic variability. σ^V^ is an extracytoplasmic function (ECF) alternative sigma factor. Alternative sigma factors replace the ‘housekeeping’ sigma factor, σ^A^, in the RNA polymerase holoenzyme and redirect it to regulons that control distinct regulatory programs. They have already been shown to display a high level of gene expression variability in *B. subtilis* (Locke et al. 2011; Young et al. 2013; *Cabeen et al. 2017; Park et al. 2018)* and the *σ*^*V*^ activation pathway is both well characterised and specific to one stress condition, which greatly simplifies analysis of its activation.

*σ*^*V*^ is the only pathway activated in response to C-type lysozyme (Ho et al. 2011; Guariglia-Oropeza and Helmann 2011; Ho and Ellermeier 2012). Lysozyme is produced by animals as part of their innate immune system, and kills bacteria by cleaving the peptidoglycan of the cell wall between the N-acetylmuramic acid residue and the β-(1,4)-linked N-acetylglucosamine (Lal and Caplan 2011). In its inactive form, σ^V^ is bound to its anti-sigma factor RsiV, a transmembrane protein (Figure 1 B). If lysozyme is present, RsiV binds to lysozyme (Hastie et al. 2014; Hastie et al. 2016) and activates a signal transduction cascade to release σ^V^. First RsiV undergoes a conformational change that allows signaling peptidases to cleave RsiV at site-1 (Lewerke et al. 2018; Hastie et al. 2014; Castro et al. 2018) (Figure 1B). *B. subtilis* has five type 1 signal peptidases, of which the two major peptidases are SipS and SipT (Tjalsma et al. 1998). Either SipS or SipT is sufficient for site-1 cleavage (Castro et al. 2018; Ho and Ellermeier 2019). The truncated RsiV can then be cleaved by RasP (a site-2 protease), which results in the release of σ^V^ (Hastie et al. 2014; Hastie et al. 2013) (Figure 1B).

Once σ^V^ is released it can bind to RNA polymerase and redirect transcription to the σ^V^ regulon, which contains genes that allow adaptation to lysozyme stress. Like most alternative sigma factors, one of σ^V^’s targets is its own operon. The *sigV* operon includes *sigV* (the gene that codes for σ^V^) and its anti-sigma factor *rsiV*, and so its activation results in ‘mixed’ positive and negative feedback loops. In addition, the operon contains *oatA*, one of the main lysozyme resistance genes, and *yrhK*, which is of unknown function (Hastie and Ellermeier 2016). OatA contributes to lysozyme adaptation by transferring an acetyl group to the C-6-OH position of N-acetylmuramic acid in the peptidoglycan, thus blocking cell wall cleavage by lysozyme (Ho et al. 2011; Bernard et al. 2011).

Although the molecular mechanisms underlying σ^V^ activation by lysozyme have been elucidated, previous work was carried out using bulk experiments that average out single cell dynamics ((Ho et al. 2011) and (Guariglia-Oropeza and Helmann 2011)) and obscure cell-to-cell heterogeneity. This heterogeneity can be crucial in identifying and distinguishing between regulatory models (Munsky et al. 2012). In this study, we used single-cell time-lapse microscopy of fluorescent reporters for σ^V^ activity to characterise σ^V^ induction dynamics in individual cells in response to lysozyme stress. We found that upon induction by sub-lethal levels of lysozyme, σ^V^ is activated heterogeneously, with some cells activating σ^V^ rapidly, whereas some cell lineages do not activate σ^V^ for multiple generations. This heterogeneity is functional, as cells that respond to an initial sublethal stress are more likely to survive a subsequent lethal stress of lysozyme. Through experiment and modelling, we found that these dynamics can be explained solely by the autoregulatory feedback of σ^V^ on its own operon. Our model predicted that the observed heterogeneity could be tuned by environmental history and by genetic perturbations, which we confirmed experimentally. The conserved sigma-anti-sigma autoregulation motif is thus a simple mechanism for bacterial populations to tune their levels of phenotypic variability.

## RESULTS

### Individual cells activate σ^V^ heterogeneously under lysozyme stress

To characterise σ^V^ activation dynamics, we first constructed a *B. subtilis* reporter strain containing a chromosomally integrated fluorescent reporter for σ^V^ activity, P_sigV_-YFP. This strain also contained a reporter for the housekeeping sigma factor σ^A^ (P_trpE_-RFP). P_trpE_-RFP expression was used as a constitutive control and to aid image analysis (Methods). We then used time-lapse microscopy to examine σ^V^ activity in individual cells grown in the mother machine microfluidic device (Wang et al. 2010). This device allows long-term tracking of hundreds of individual cells trapped at the ends of channels. It also allows fast switching between media conditions during imaging (Methods). We first measured σ^V^ expression dynamics in response to a step change to a sub-lethal concentration of 1 µg/ml lysozyme (Figure 1C & D,Figure S1 & Figure S2 and Movie S1). The first cells begin to respond to the lysozyme stress with raised P_sigV_-YFP levels within 2 frames (20 minutes), suggesting rapid activation given the ∼15 minute maturation time of the YFP reporter protein (Figure S3). However, our time-lapse movies revealed that P_sigV_-YFP was heterogeneously activated at the single-cell level (Figure 1C). While 10% of cells reached the half maximum of σ^V^ activity within 70 min of being exposed to lysozyme, it took 200 min, (approximately 4 generations) for 90% of all cells to activate σ^V^ (Figure 1D and Figure S4B).

To test whether the observed heterogeneous activation of σ^V^ in response to lysozyme was due to the growth conditions in the mother machine, we investigated σ^V^ activation dynamics in liquid culture (Figure S5) and using an alternative microfluidic device (Figure S6). Both conditions showed similar heterogeneous σ^V^ activation to that observed in the mother machine. Therefore the observed heterogeneity in P_sigV_-YFP is independent of the experimental setup. Previous work has shown that heterogeneous gene expression could be due to intrinsic noise in the expression of the P_sigV_-YFP reporter (Elowitz et al. 2002), rather than due to heterogeneous σ^v^ activity. To test whether the observed heterogeneity was due to the intrinsic variability of the P_sigV_-YFP reporter we constructed a strain containing chromosomally integrated P_sigV_-YFP and P_sigV_-CFP reporters and exposed it to 1 µg/ml lysozyme in liquid culture. In these snapshot experiments, the expression of P_sigV_-YFP and P_sigV_-CFP was highly correlated (R^2^ = 0.85) (Figure S7). Thus the observed heterogeneity in P_sigV_-YFP reflects changes in σ^v^ activity and not intrinsic variability of the P_sigV_-YFP promoter.

Next, we asked whether the observed heterogeneity in P_sigV_-YFP is modulated by the level of lysozyme applied. We examined P_sigV_-YFP expression after application of 0.5, 1, 2, and 4 µg/ml lysozyme. These values were all below the previously reported minimal growth inhibitory concentration of 6.25 µg/ml (Ho et al. 2011). To measure the distribution of σ^V^ activation times, for each time point we calculated the fraction of cells that had crossed the half-maximum of their respective final σ^V^ value (meaning the cell had activated σ^V^). We found that when increasing the lysozyme concentration from 0.5 µg/ml to 4 µg/ml the heterogeneity in σ^V^ activity was reduced (Figure 1E, Figure S2 and Figure S8). The time for 90% of cells to activate their σ^V^ pathway decreased from 300 min (approximately 6 generations) for 0.5 µg/ml to 100 min (approximately 2 generations) for 4 µg/ml lysozyme (Figure S4). At the same time, the fold change in induction between the unstressed σ^V^ activity and the steady state σ^V^ activity under lysozyme stress increased from ∼190 for 0.5 μg/ml to ∼520 for 4 μg/ml (Figure 1F and Figure S1). We also observed that under a 4 µg/ml concentration of lysozyme some cells (8% and 21% in two different repeat experiments) appeared sick and were wider than usual cells. These cells also overshot their new σ^V^ activity steady state before relaxing to it. We removed these cells from our analysis (Figure S9), although including them did not affect our results (Figure S10). Taken together, our results reveal that the level of phenotypic variability in *sigV* expression is tuned by stress levels, with P_sigV_-YFP heterogeneity reduced as lysozyme levels increase.

We next asked whether differences in individual cell states before applying a lysozyme stress could predict how a cell responds to the stress. There was no correlation between either growth rate or cell size before lysozyme application and response time after lysozyme application (Figure S11 and Figure S12). Alternative sigma factors in *B. subtilis* display gene expression variability even in the absence of stress (Locke et al. 2011; Park et al. 2018). We found that, in the absence of stress, the coefficient of variation of P_sigV_-YFP was over 4 times higher than that of the constitutive control (0.65±0.29 versus 0.14±0.05) (Figure S13). Cells which had elevated levels of P_sigV_-YFP before the addition of stress were more likely to activate σ^V^ instantaneously (Figure S14 and Figure S15). However, some cells with low P_sigV_-YFP levels could also activate σ^V^ instantaneously (Figure S14 and Figure S15). Therefore, while noise in P_sigV_-YFP expression before stress contributes to the observed heterogeneity in σ^V^ activation after the addition of lysozyme, it is not the only cause.

Next, we examined whether activating σ^V^ had any effect on the survival against future lethal concentrations of lysozyme. Cells were first exposed to a priming stress of 1 μg/ml of lysozyme for 30 min, which induced σ^V^ heterogeneously. Thereafter cells were exposed to 20 μg/ml of lysozyme for 20 min before removing lysozyme from the media (Figure 2A). Surviving cells were defined as cells which had not lysed at the end of the movie, 280 min after the addition of 20 μg/ml of lysozyme. We found that the short exposure to sub-lethal concentrations of lysozyme (during the priming stress) improved survival to subsequent lethal stress levels (Figure 2B) from 0% (Figure S16A) to 12 ± 5% survivors (Figure S16B). Surviving cells had on average 1.57 ± 0.33 fold higher P_sigV_-YFP levels than perishing cells immediately before the application of the lethal concentration of lysozyme (Figure 2C).

**Figure 2.**
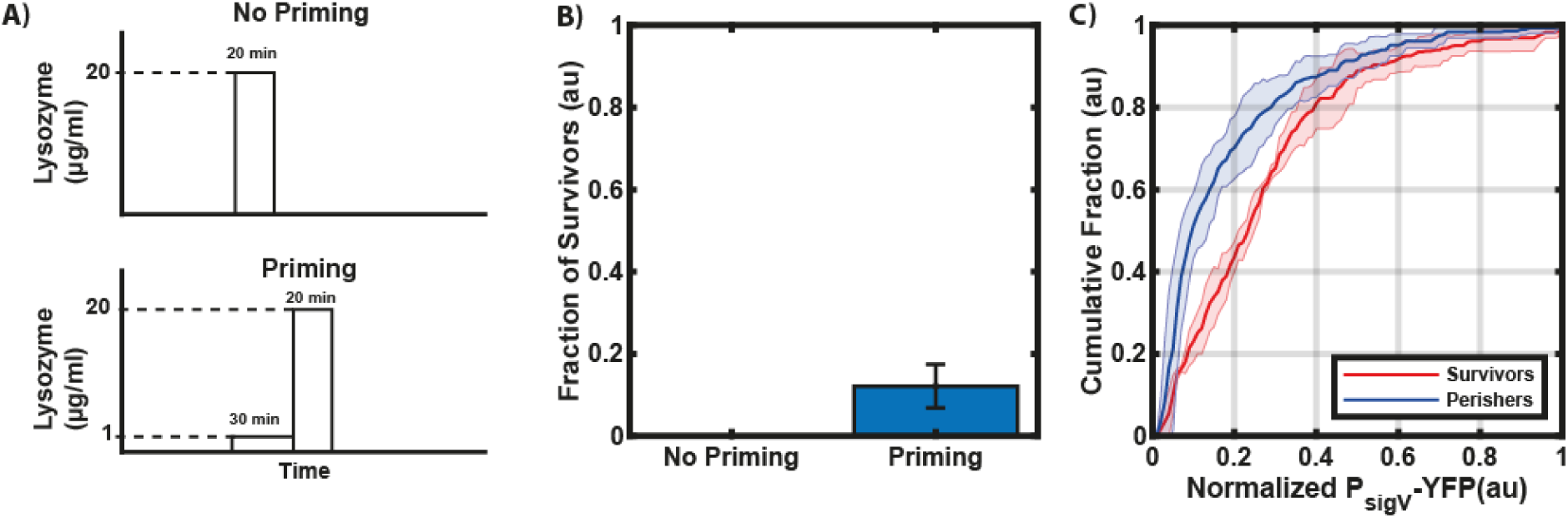
Rapid activation of σ^V^ after a first stress application increases survival after a second higher stress. (A) Schematic of lysozyme application. Cells are either exposed directly to a high concentration of lysozyme (20 µg/ml) for 20 min (top) or exposed first to a short (30 min) lysozyme priming stress (1 µg/ml) before exposure to the higher concentration (bottom). (B) A priming (30 min) stress of 1 µg/ml lysozyme followed by the high lysozyme stress (20 µg/ml) improves survival. (C) Surviving cells have higher P_sigV_-YFP levels after the initial priming stress (1 µg/ml) than perishing cells. The cumulative distributions were normalized by their maximum σ^V^ activity and baseline subtracted. Each line is the mean of experiments from n=4 biological repeats. The shaded areas correspond to the mean ± SD. For more information on the number of repeats and cell numbers please see the supplementary text.

### The heterogeneous activation of P_sigV_-YFP is solely due to σ^V^ and its anti-sigma factor RsiV

We next attempted to understand how the single-cell activation dynamics of the σ^V^ pathway are generated. First, to test whether the heterogeneity that we observed in σ^V^ activation times was due to some cells activating a different stress response pathway, we analysed the genome–wide transcriptional response of cells to 1 μg/ml lysozyme in the wild type and in the Δ*sigV* knockout. As previously reported, only the *sigV* operon was induced by lysozyme (Guariglia-Oropeza and Helmann 2011) in the wild type (Figure 3B). The Δ*sigV* strain did not show any induction (Figure 3C). This suggests that no other pathways are induced by lysozyme. We therefore hypothesized that the observed P_sigV_-YFP heterogeneity is due to the σ^V^ circuit itself (Ho et al. 2011; Hastie et al. 2013; Hastie et al. 2014).

**Figure 3.**
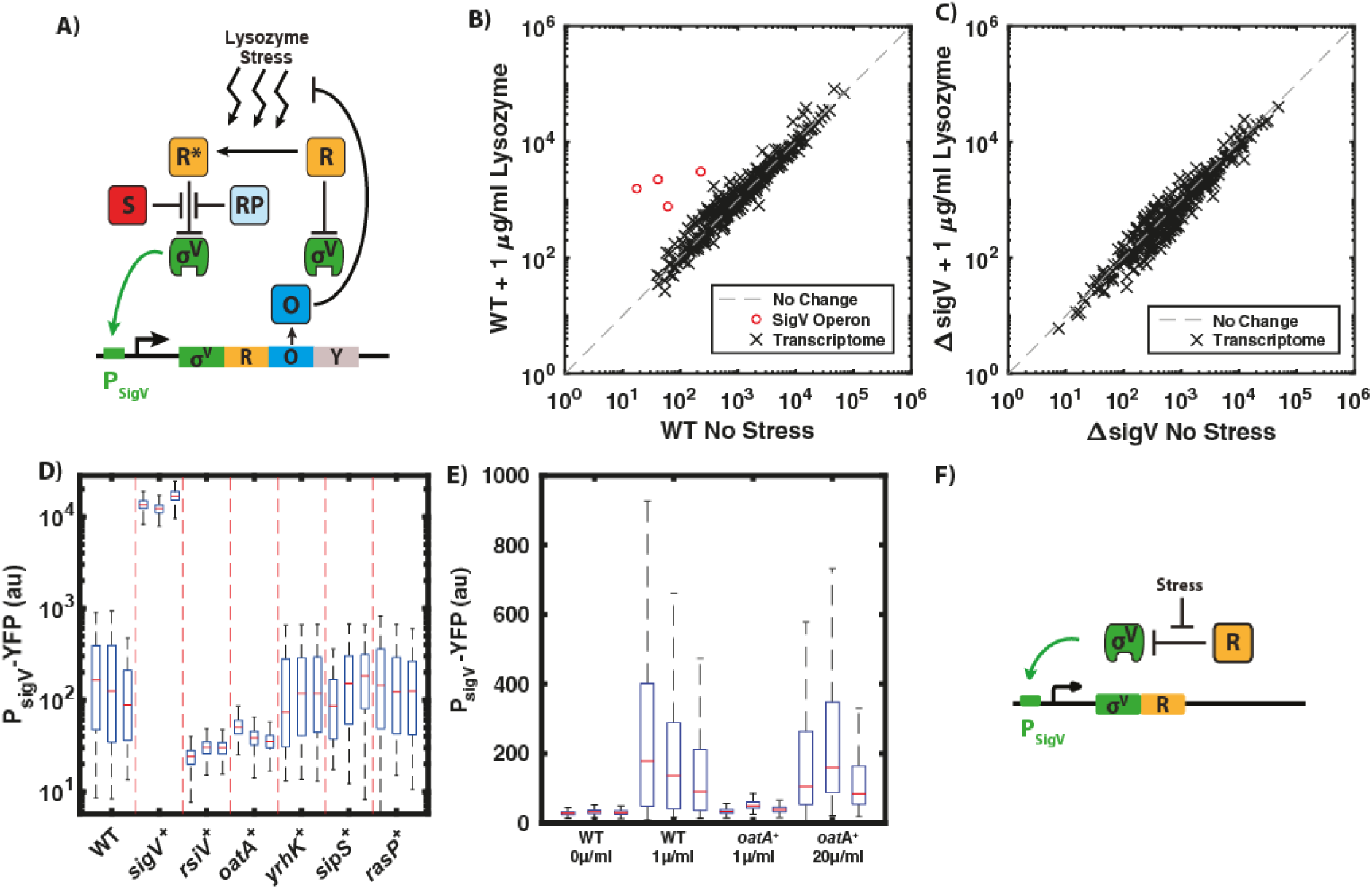
The observed σ^V^ heterogeneity can be explained by a simplified σ^V^ circuit. (A) Schematic of the σ^V^ circuit. In the figure R (orange) is the anti-sigma factor RsiV, R* (orange) is RsiV bound to lysozyme, S (red) is signaling peptidase and RP (light blue) is RasP the site-2 protease. For more information on the activation mechanism see Figure 1B. (B &C) RNA-seq screen of WT(JLB130) and Δ*sigV(*JLB154) strains, showing quantification of the expression of individual genes in the presence and absence of stress. (B) Only the *sigV* operon is activated in response to lysozyme stress in WT (JLB130), as previously reported (Guariglia-Oropeza and Helmann 2011). (C) No genes were significantly upregulated in Δ*sigV* (JLB154) by lysozyme stress. (D) Effect of the overexpression of individual components of the σ^V^ pathway (WT; n=8, *sigV+;* n=4, *rsiV+*; n=3 *oatA+;* n=4, *yrhK+;* n=3, *sipS+;* n=3 and *rasP+;* n=3) on the level of sigV activation. Only overexpression of *sigV, rsiV* or *oatA* changed the observed dynamics compared to WT. Only three randomly selected boxplots are shown for circuit components where n>3. (E) Overexpressing *oatA* shuts off σ^V^ activation. Increasing stress levels from 1 µg/ml to 10 µg/ml recaptures the WT σ^V^ activation dynamics. n>=3 for all data shown. Only three randomly selected boxplots are shown for circuit components where n>3. (F) Schematic of simplified σ^V^ circuit with only σ^V^ and RsiV, where σ^V^ (green) activates its own expression and that of its anti-sigma RsiV (orange, R).

To test which components of the σ^V^ circuit play a role in the heterogeneous activation of σ^V^, we constructed IPTG inducible overexpression constructs of all genes in the *sigV* operon (*sigV, rsiV, oatA, yrhK*) and in the σ^V^ circuit (*sipS, rasP*) (Figure 3A). We then investigated σ^V^ activation dynamics under 1 μg/ml lysozyme after full induction of each circuit component. Single-cell snapshots of P_sigV_-YFP expression revealed that overexpressing either *yrhK, sipS*, or *rasP* did not have any effect on the heterogeneous activation of σ^V^ (Figure 3D and Figure S17). Overexpression of *sigV* itself removed the heterogeneity in P_sigV_-YFP as all cells activated even before the addition of lysozyme (Figure S17). Conversely, overexpression of *rsiV* caused cells not to express P_sigV_-YFP at all (Figure 3D and Figure S17). Finally, overexpressing *oatA*, which blocks lysozyme cleavage of the peptidoglycan, shuts off the activation of σ^V^ (Figure 3D and Figure S17). However, when we increased the lysozyme concentration to 10 μg/ml in the presence of *oatA* overexpression the heterogeneous expression of P_sigV_-YFP reappeared (Figure 3E), suggesting oatA is not responsible for the heterogeneous activation dynamics. These findings suggest that the heterogeneity in σ^V^ activation only depends on σ^V^ and its anti-sigma factor RsiV (Figure 3F).

### Mathematical modelling reveals that P_sigV_-YFP expression dynamics can be explained by the ‘mixed’ σ^V^ autoregulatory feedback loop

Based on our transcriptome and overexpression analysis we constructed a model of a simplified *sigV* operon consisting of *sigV* and its anti-sigma factor *rsiV*, with positive autoregulation of the operon by active σ^V^ (Figure 3F). We did not include *oatA*, as it is not required for the heterogeneous activation of σ^V^ (Figure 3F). RsiV binds σ^V^ to form an inactive complex σ^V^-RsiV. On stress application, RsiV is degraded and σ^V^ is free to activate the operon. The model was implemented using mass action kinetics, with hill equation terms for production rates. Chemical Langevin equations were used to model the intrinsic noise of the pathway, creating a system of stochastic differential equations (See methods for further details, (Gillespie 2000)).

Using a coarse-grained parameter search, we were able to find parameters that capture the heterogeneous σ^V^ activation in response to lysozyme stress (Figure 4A and Figure S18). We were also able to recreate several aspects of the experimental data, including the dependency of induction time (Figure 4B) and steady state levels of *sigV* expression on increasing levels of lysozyme stress (Figure 4C). In the absence of lysozyme stress, the system is monostable and only a state with low σ^V^ activity is accessible (Figure S19). Application of stress introduces a second steady state of high σ^V^ activity, whilst also making the low sigV state only quasi-stable (Figure S19 B). Noise in sigV expression causes a stochastic switch to the high σ^V^ state. The stochastic nature of the switch generates the broad distribution of induction times observed (Figure S18).

**Figure 4.**
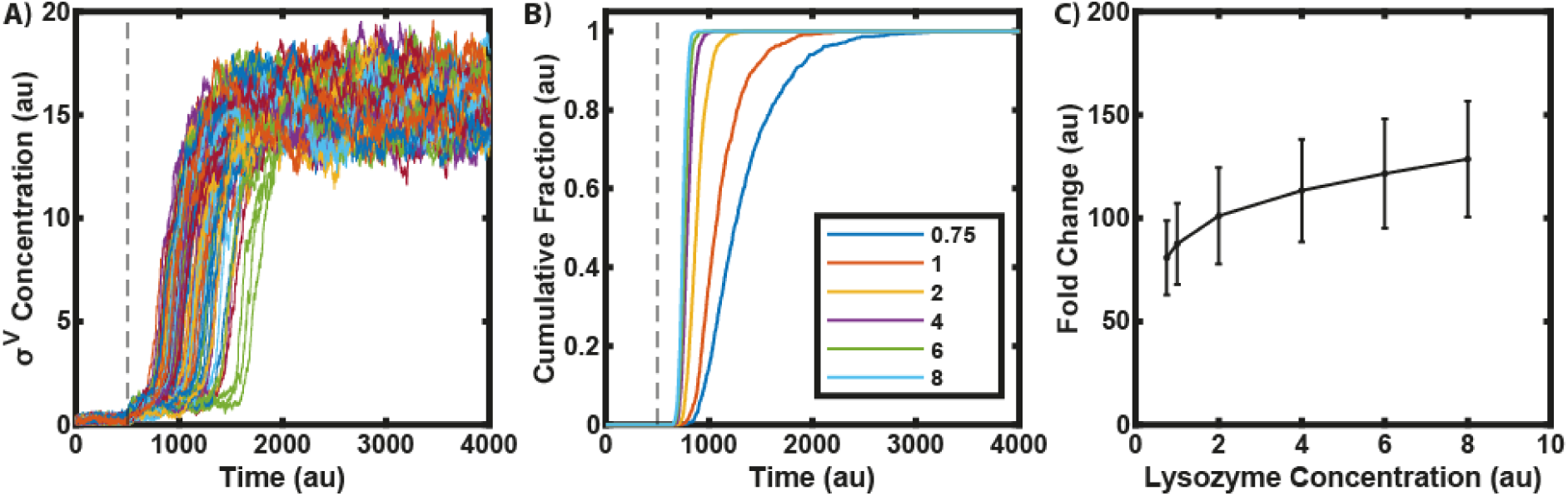
A model of the simplified σ^V^ circuit captures experimental results. (A) Simulations display heterogeneous σ^V^ activation after lysozyme stress (N=100). Dotted vertical line marks point of stress addition in simulation. (B) In simulations cells activate σ^V^ faster as stress levels are increased, as observed experimentally (N=1000 for each stress level). (C) With increasing stress levels, the steady state level of σ^V^ activity increases, as observed experimentally. Error bars correspond to the mean ± SD (N=1000 for each lysozyme stress level).

### Phenotypic variability is tuned by doubling copy numbers of *sigV* operon genes

To test the assumptions of our model, we predicted the effect on σ^V^ activation dynamics of introducing a second copy of each component of the *sigV* operon (*sigV, rsiV*, or both *sigV* and *rsiV*), with each component driven by the *sigV* promoter. Our model predicted that these perturbations would modulate the heterogeneity of the system’s response to lysozyme stress. A second copy of *rsiV* meant that larger fluctuations in *sigV* expression were required to kick the system into the high σ^V^ expression state (Figure 5B and Figure S20). This increased the cell to cell variability in response times to lysozyme stress, as compared to the WT in the simulations (Figure S20). Conversely, with a second copy of *sigV*, or a second copy of both *sigV and rsiV*, smaller fluctuations in sigV expression are required to kick the system into the high σ^V^ expression state. Cell to cell variability in response times to lysozyme therefore decreases in the simulations (Figure 5C and Figure S20). In fact, a second copy of *sigV* caused an increase in the levels of σ^V^ expression even before the addition of lysozyme in the simulations (Figure S20).

**Figure 5.**
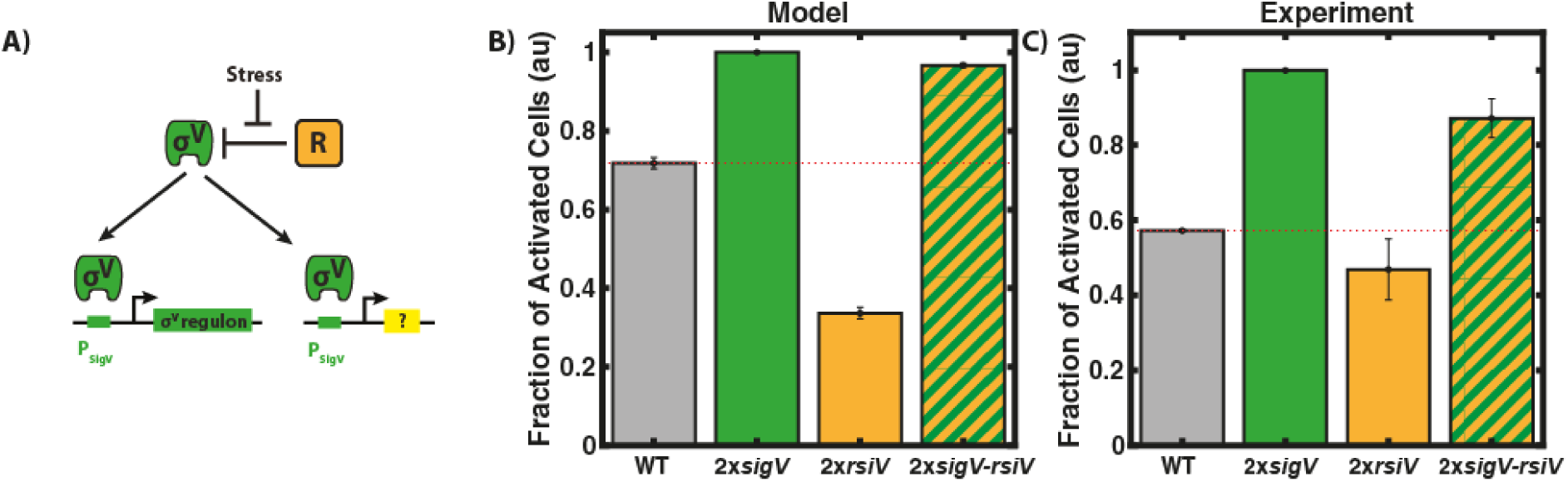
In both simulations and experiment, phenotypic variability is tuned by additional copies of components of the σ^V^ circuit. (A) Schematic of stochastic tuning with an additional copy of the σ^V^ circuit. (B) Model simulations predict that a second copy of P_sigV_-sigV (2x*sigV*) or P_sigV_-sigVrsiV (2x*sigV-rsiV*) makes the σ^V^ response to lysozyme homogenous while a second copy of P_sigV_-rsiV (2x*rsiV*) should increase the heterogeneity. Bars represent the fraction of cells which have activated σ^V^ 30 min after treatment with lysozyme. Cells which had activated σ^V^ were defined as cells which had passed a threshold of the WT mean before lysozyme application plus six standard deviations. Error bars correspond to mean (N=999) ± SD (1000 bootstraps). (C) As predicted, adding a second copy of P_sigV_-sigV (2x*sigV*) or P_sigV_-sigVrsiV (2x*sigV-rsiV*) makes the σ^V^ response to 1 µg/ml lysozyme homogenous in experiments, while a second copy of P_sigV_-rsiV (2x*rsiV*) increased the heterogeneity. Each bar plot is the average of three biological repeats. The error bars correspond to the mean ± SD.

To test these predictions, we constructed strains containing a second copy of either *sigV, rsiV* or *sigV-rsiV* driven by the *sigV* promoter. As predicted by our model, snapshots of *sigV* expression after induction by 1 μg/ml of lysozyme revealed that a second copy of *sigV* or *sigV-rsiV* reduced the observed heterogeneity in σ^V^ activation, whilst a second copy of r*siV* caused an increase in the heterogeneity in σ^V^ activation (Figure 5C). Finally, adding a second copy of *sigV* alone caused the activation of σ^V^ even in the absence of stress, as predicted. These results validate our model assumptions, and also show that the heterogeneity in *sigV* expression is easily tuned by simple genetic perturbations.

### The heterogeneous activation dynamics of *sigV* are dependent on environmental history

Given that the σ^V^ activation dynamics appear sensitive to the baseline levels of σ^V^, RsiV, and the σ^V^-RsiV complex, any perturbations to these levels should affect σ^V^ activation times. We hypothesized that cells should have elevated levels of *sigV* operon components after a lysozyme stress is removed due to the recent high expression of the operon components. The elevated levels of σ^V^ stored in stabilized σ^V^-RsiV complexes should cause all cells to respond immediately to a subsequent reapplication of lysozyme. We investigated this hypothesis in our model. We simulated the application of lysozyme stress, which was then removed for the equivalent duration of several cell cycles The same level of stress was then reapplied and the heterogeneity in σ^V^ activation disappeared (Figure 6A and Figure S22). However, as the delay before the second application of lysozyme was increased (allowing system components to relax to pre-stress levels), the heterogeneity gradually returned (Figure 6B&C and Figure S22). Thus our simulations predict the existence of a temporary molecular memory of the environmental history.

**Figure 6.**
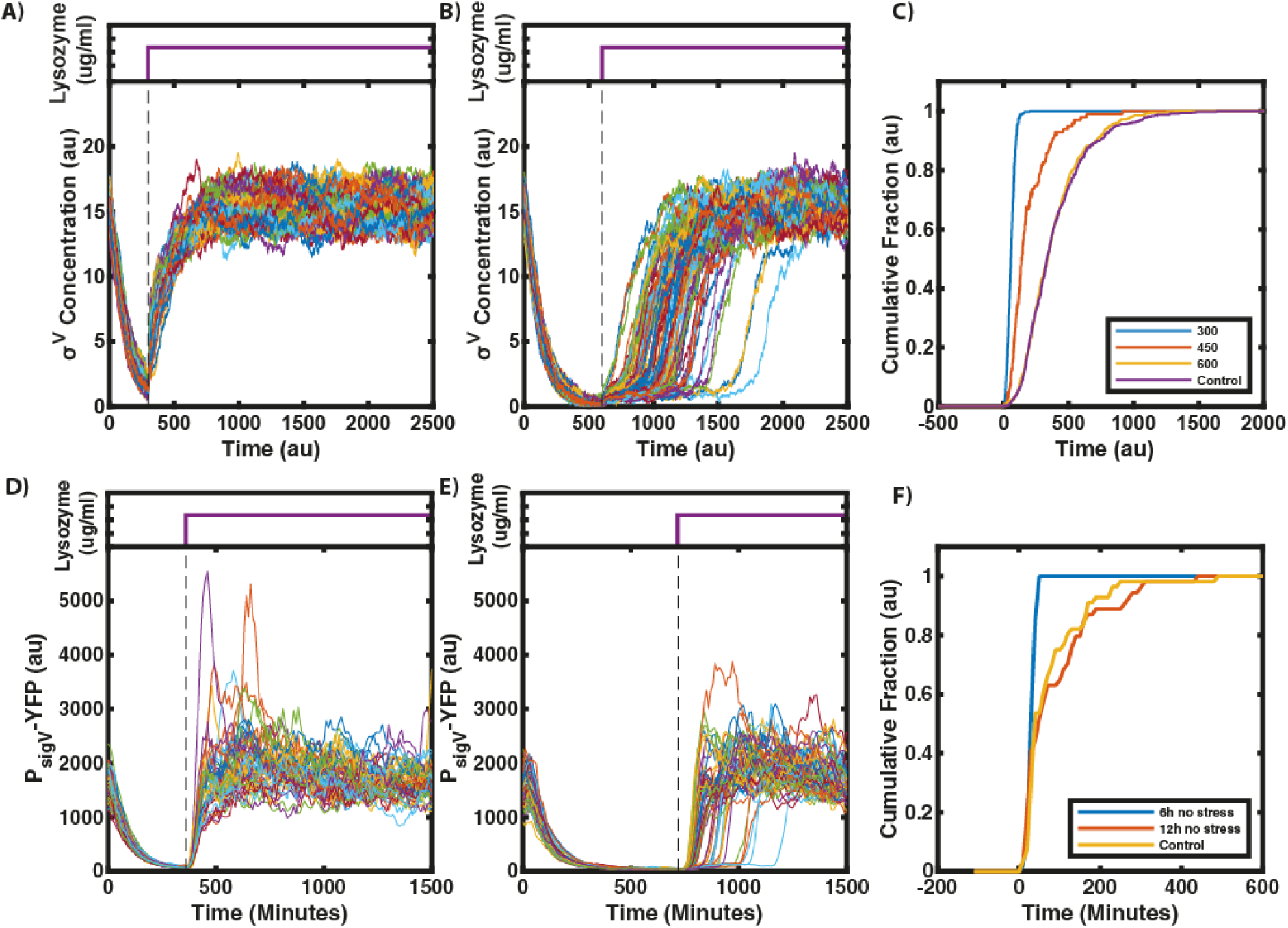
Modelling predicts a memory of previous stress. Stress is removed at time 0 and reapplied at a time indicated by the dashed line, lysozyme concentrations are indicated by the purple traces above time course plots. (A) Our model predicts that after a break in stress, σ^V^ activation is homogenous (N=100). (B&C) With increasing intervals between stresses, response heterogeneity reappears in the model, indicating a loss of memory. (C) Each line represents the cumulative fraction of cells (N=100) with σ^V^ concentration values that are higher than the half maximum of their final values (representing cells that have activated). If the interval between stress is increased long enough, (600 au), the heterogeneity is the same as if the cells had not experienced a prior lysozyme stress (control). (D) This memory effect can be reproduced in the experiments. For short intervals between stress applications, cells respond immediately and homogeneously. Each line corresponds to one of 48 single cell traces. (E & F) For very long intervals between stress (12h), the heterogeneity is the same as if the cells had not experienced a prior lysozyme stress (control). N>50 single cell traces.

To test whether the σ^V^ circuit exhibits such a memory, we grew cells in the mother machine device under 1 μg/ml lysozyme stress before removing the stress for 6h (approximately 7 cell cycles). All cells stopped activating σ^V^ as soon as lysozyme was removed, as indicated by the decay in YFP fluorescence (Figure 6 D and Movie S2). The decay in fluorescence was tightly synchronized across the population and YFP decayed with a half-life time of 51 ± 1 min, which was similar to the cell cycle time of 51 ± 13 min. When reapplying a 1 µg/ml lysozyme stress after 6h all cells responded instantaneously. As predicted by the model, the heterogeneity in σ^V^ activation was lost (Figure 6D,Figure S25A and Movie S2). Also as predicted by the model, increasing the duration of the break in stress to 12h resulted in heterogeneous activation dynamics similar to those observed in response to the first application of lysozyme stress (Figure 6E&F, Figure S25 B&C and Movie S3). This return to a heterogeneous response reflects the system’s loss of memory - through reduction in the *sigV* operon component concentrations to pre-stress exposure levels. These results held for two different versions of the mother machine microfluidic device (Figure S26). Taken together, our results show how the simple autoregulatory σ^V^ circuit can generate heterogeneous activation dynamics that can be tuned by stress level, genetic perturbations, and the environmental history of the cell.

## DISCUSSION

Here, we report a general mechanism for a bacterial population to tune its phenotypic variability based on stress levels, genetic architecture, and environmental history. Using quantitative single-cell microscopy and microfluidics we found that the activation of the alternative sigma factor σ^V^ in response to lysozyme stress was heterogeneous. While some cells activated their σ^V^ pathway immediately, others could take up to six generations to activate their σ^V^ pathway. The observed phenotypic variability plays a functional role, as cells that respond to a sub-lethal stress were more likely to survive a subsequent higher stress application (Figure 2). Through experiments and modelling we found that this heterogeneity could be understood by the ‘mixed’ positive and negative feedback of σ^V^ activating both itself and its anti-sigma factor RsiV. Alternative sigma factors are a common regulatory system in prokaryotes, often controlling stress response and virulence pathways. The ‘mixed’ feedback loop is also a common motif, suggesting that this motif can be a general mechanism for bacterial populations to tune their phenotypic variability.

Our modelling and experiments found that recent exposure to lysozyme stress is remembered, even though the system turns off after removal of lysozyme stress. The key to this behaviour appears to be the ‘mixed’ feedback loop, as it allows amplified levels of inactive σ^V^-RsiV complexes in each cell after stress. These complexes can be cleaved by a subsequent addition of lysozyme, releasing σ^V^ to activate its operon. Similar memories of previous stress have been observed in bacterial systems, although typically not to modulate phenotypic heterogeneity. For example other pathways such as the heat stress response in *B. subtilis* (Runde et al. 2014) or the oxidative stress response in yeast (Kelley and Ideker 2009) have been shown to have a memory of past conditions. Often this memory is facilitated through an autoregulatory positive feedback loop that can lock the cell into an ON state that is heritable through cell divisions (Lambert and Kussell 2014; Novick and Weiner 1957) (Zacharioudakis et al. 2007; Acar et al. 2005; Hashimoto et al. 2013; Biggar and Crabtree 2001). However, it is difficult for a single positive feedback loop to allow the system to be OFF but primed for future stress, as we find to be the case for the ‘mixed’ feedback loop in the σ^V^ pathway. In the future, it will be important to investigate whether the ‘mixed’ feedback loop also tunes the levels of phenotypic diversity by environmental history in other systems. Interestingly, computational studies have proposed that a ‘mixed’ feedback loop structure in the MAR operon in *E. coli* allows the tuning of the fraction of cells prepared to survive antibiotic exposure (Garcia-Bernardo and Dunlop 2013).

We have found that a combination of noise in gene expression and a mixed feedback loop can generate tunable phenotypic diversity in an alternative sigma factor circuit. Going forward, it will be important to observe whether alternative sigma factors more generally are used as a mechanism for bacterial populations to tune phenotypic diversity. This is particularly the case given that alternative sigma factors often control pathways critical to pathogenicity and resistance to antibiotics. Noise in sigma factor activity has already been observed in multiple alternative sigma factor circuits in *B. subtilis* (Locke et al. 2011; Park et *al. 2018*), as well as for the general stress response sigma factor in *E. coli* (Patange et al. *2018*). Additionally, an alternative sigma factor plays a role in generating phenotypic diversity in Mycobacteria (Ghosh et al. 2011; Sureka et al. 2008). Two other obvious candidates for further study are the pathogens *Enterococcus faecalis* and *Clostridioides difficile* (Ho and Ellermeier 2019), which also have a σ^V^ pathway that is responsive to lysozyme.

## MATERIAL AND METHODS

### Strains and Media

All strains are derivatives of the PY79 background strain (Table S1). Deletions were generated by replacing genes of interest with an antibiotic resistance cassette by recombination of a linear DNA fragment homologous to the region of interest. All strains had a Δ*ytvA*::neo deletion insertion. YtvA is a blue light sensor and it was deleted to avoid any activation of the general stress response pathway σ^B^ by the microscope illumination (Locke et al. 2011; Gaidenko et al. 2006). For strains used in mother machine experiments, cells were made immotile by inserting a Δ*haG*:erm deletion cassette in order to improve the loading into the microfluidic device. Additionally, all strains contained a house keeping σ^A^ promoter-driven mCherry for segmentation and as a constitutive control. (Locke et al. 2011).

Cells were routinely grown in Spizizen’s Minimal Media (SMM) (Spizizen 1958). It contained 50 µg/ml Tryptophan as an amino acid source and 0.5% glucose as a carbon source. Cultures were started from frozen stock in SMM and grown overnight at 30°C to an OD between 0.3 and 0.8. The overnight cultures were resuspended to an OD of 0.01 and regrown to an OD of 0.1 at 37°C.

### Plasmids

*E. coli* strain DH5α was used to clone all plasmids. The cloning was done with a combination of non-ligase dependent cloning and standard molecular cloning techniques using Clonetech In-Fusion Advantage PCR Cloning kits. Plasmids were chromosomally integrated into the PY79 background via double crossover using standard techniques. The list below provides a description of the used plasmids, with details on selection marker and integration position/cassette given at the beginning. Note that all plasmids below replicate in *E. coli* but not in *B. subtilis*.

#### 1. ppsB::P trpE-mCherry ErmR

This plasmid was used to provide uniform expression of mCherry from a σ^A^-dependent promoter, enabling automatic image segmentation (cell identification) in time-lapse movie analysis. A minimal σ^A^ promoter from the trpE gene was cloned into a vector with ppsB homology regions (Locke et al. 2011). The original integration vector was a gift from A. Eldar (Eldar et al. 2009).

#### 2. sacA::P sigV -yfp CmR

The target promoter of sigV was cloned into the EcoRI/BamHI sites of AEC12 (Eldar et al. 2009) (gift from M. Elowitz, CalTech).

#### 3. amyE::P sigV -mturq SpectR

The target promoter of sigV was cloned into the EcoRI/Nhe1 sites of plasmid amyE ::3XCFP Spect R (Locke et al., 2011). Target promoter sequences are described below.

#### 4. amyE::P hyperspank -? SpectR

The coding region of (where ? is sigV, rsiV, yrhK, oatA, rasP, or sipS) along with a 5’ transcriptional terminator, was cloned downstream of the hyperspank IPTG-inducible promoter in plasmid pDR-111 (gift from D. Rudner, Harvard).

#### 5. amyE::P sigV-? SpectR

The coding region of (where ? is sigV, rsiV, sigVrsiV) along with a 5’ transcriptional terminator, was cloned downstream of the sigV promoter.

### Microscopy

A Nikon inverted Ti-E microscope (Nikon, Amsterdam, Netherlands) with a Nikon Perfect Focus System (PFS) hardware auto-focus was used. All images were acquired using either a Photometrics CoolSnap HQ2 CCD camera (Photometrics, Tucson, AZ, USA) or a Photometrics Prime sCMOS camera (Photometrics, Tucson, AZ, USA). The microscope stage was enclosed in an incubator (Solent Scientific, Segensworth, UK) which was set to 37°C for all experiments. Illumination was provided by a LED lamp (CoolLED, Andover, UK). Epifluoresence was provided by a Lumencore Solar II light engine (Lumencore, Beaverton, OR, USA). Chroma filters (Chroma, Bellows Falls, USA) #41027 for the RFP channel, #49003 for the YFP channel and #49001 for the CFP channel were used. All experiments were done with a phase 100x Plan Apo (NA 1.4) objective (Nikon, Amsterdam, Netherlands). Metamorph (Molecular Device, Sunnyvale, CA, USA) controlled the camera, the motorized stage (Nikon, Amsterdam, Netherlands) and the microscope.

#### Snapshots

Agarose pads were made by pipetting 1 ml of 1.5 % (wt/vol) low melt agarose (Merck, Darmstadt, Germany) dissolved in PBS onto a 22 mm^2^ cover glass slide. Immediately after pipetting, another cover glass slide was placed on top of the agarose creating a “sandwich” with the agarose in the middle. Once the agarose had solidified, the top cover glass was removed and the agarose was cut into squares of approximately 5 mm × 5 mm with a scalpel.

Once the cultures had reached an OD of 0.1, they were split into aliquots of 1 ml to which different concentrations of lysozyme from hen eggs white (Sigma Aldrich, St. Louis, MO, USA) were added. The aliquots were then incubated at 37°C for 30 min. For experiments with IPTG inducible strains, 1 mM of IPTG was added to the aliquots for 60 min before addition of lysozyme, in order to allow for full induction of the promoter before stress. Different concentrations of lysozyme were then applied for 30 min before snapshot measurements. For snapshots, 2 µl of cell culture were pipetted onto the pads, which were left to dry and then laid face down onto a cover glass-bottom dish (#HBSt-5040, WillCO-dish, Amsterdam, Netherlands). The glass dish was sealed with parafilm and put under the microscope to image. Single-cell data was extracted using custom MATLAB (Mathworks, Natik, USA) scripts based on the Schnitzcells package (Young et al. 2011).

#### CellAsic Experiments

Cultures were grown overnight to an OD of 0.1, and then resuspended to an OD of 0.001 for loading into CellAsic B04 microfluidic chips (Merck, Darmstadt, Germany). Cells were loaded into the microfluidic chips with a pressure of 4 to 6.5 psi for 2 to 4 s. The following lysozyme concentrations were investigated with the CellAsic setup: 0 µg/ml, 0.5 µg/ml, 1 µg/ml and 4 µg/ml. Fresh media was perfused into the chip with a pressure of 1 psi. After 60 min of growth in standard SMM, the media was switched to media containing lysozyme. Cells were imaged at regular intervals (every 10 min) and the acquired movies were analysed with the standard Schnitzcells package for MATLAB (Young et al. 2011).

### Microfluidics

#### Wafer Fabrication

We used two different microfluidic designs of the mother machine

1. All the mother machine data in this paper were acquired with a microfluidic design based on the original mother machine (Wang et al. 2010) unless stated otherwise. As a substrate for the PDMS chip fabrication we used an epoxy master which was a kind gift from the Jun lab at the University of California, San Diego, USA (Taheri-Araghi et al. 2015).
2. For data shown in Figure S26 a different mother machine design was used. It was based on a mother machine design first described by Norman et al. (Norman et al. 2013). The main difference to the Jun lab design is that the channels in which cells grew were longer and that they have a shallow side channel for better perfusion of media to the cells. As a substrate for the PDMS chip fabrication we used an epoxy master which was a kind gift from the Elowitz lab at the California Institute of Technology, Pasadena, USA (Park et al. 2018)

#### Soft Lithography

Sylgard 184 polydimethylsiloxane (PDMS) from Dow Corning (Midland, MI, USA) was used to replica-mold the microfluidic devices. A 10:1 (base:curing agent) PDMS mixture was cast onto the epoxy master and cured overnight at 65°C. In the next morning, the chips were cut out and the inlets were punched using a 0.75 mm biopsy puncher (#504529, WPI, Sarasota, FL, USA). To remove any uncured PDMS from the chips, these were chemically treated in a pentane (Sigma Aldrich, St. Louis, Mo, USA) bath for 90 min, and then washed twice for 90 min in acetone (Sigma Aldrich, St. Louis, Mo, USA). The chips were then left to dry overnight in the fume hood and were ready to use the next day (Gruenberger et al. 2013).

### Experimental Preparation

The chips were plasma bonded to a glass bottom dish (HBSt-5040, Willco Wells, Amsterdam Netherlands) by treating the chip and the glass dish surfaces with a plasma (Femto Plasma System, Diener, Ebhausen, Germany) for 12 s at power of 30 W and a pressure of 0.35 mbar. The chips were then put into the oven for 10 min at 65°C, in order to strengthen the bonding. In the meantime, a bovine serum albumin (BSA) passivation buffer was prepared at a concentration of 20 mg/ml in SMM. The chips were then passivated with this buffer for 1 h at 37°C.

Cells were grown overnight as for the other movie experiments. 10 ml of the regrown cultures with an OD of 0.1 were spun down for 10 min at 4000 rpm and resuspended in 0.1 ml to a final OD of 10. This cell suspension was loaded into the growth channels by spinning the chip for 5 min at 3000 rpm with a spin coater (Polos SPIN150i, SPS, Harlem, Netherland).

#### Experimental Setup

Perfusion was provided by a Fluigent pressure pump setup. Briefly, reservoirs were pressurized using a computer controllable pressure controller (MFCS-EZ, 1bar, Villejuif, Fluigent, France). A rotary selection valve (M-Switch, Fluigent, Villejuif, France) allowed for switching between different reservoirs. The flow rate was set to 1 ml/h and by using a flow sensor (M-Flow Sensor, Fluigent, Villejuif, France) which feeds back onto the pressure controller.

For the experiments on the memory in the σ^V^ response, syringe pumps (Legato 200, KDScientific, Holliston, USA) were used. Switching was provided by an electronic switching valve (MXP9960, Rheodyne, Lake Forest, USA). The flow rate remained at 1 ml/h. The analysis of all movies was done using a modified version of Schnitzcells (Young et al. 2011) optimized for mother machine experiments.

#### Memory Experiments

For all shown memory experiments, the JLB221 strain (ΔespH) was used instead of JLB130, unless stated otherwise. This was done in order to prevent clogging of the microfluidic chip during these long term experiments due to the production of extracellular matrix. The two strains had the same P_sigV_-YFP dynamics in response to lysozyme (Figure S23 & Figure S24).

Cells were inoculated in SMM and grown overnight to an OD of 0.3 to 0.8. In the morning, the cells were resuspended to an OD of 0.01 in SMM + 1 μg/ml lysozyme and grown to an OD of 0.1. Cells were then loaded as described in the “experimental preparation” section. In the mother machine the cells were grown for 4h with 1 μg/ml lysozyme before removing the stress. The stress was then reapplied after 6h or 12h.

### Data Analysis

#### Cumulative Activation Time

The activation time was calculated as the time between switching from SMM to lysozyme and the time point when then P_sigV_-YFP passed its half maximum (Figure S1). The cumulative fraction was then calculated as the cumulative fraction of cells with P_sigV_-YFP values higher than the half maximum of their final values (representing cells that have activated) (Figure S1 and Figure S8).

#### Removal of Overshooting Cells

For 4 μg/ml lysozyme the P_sigV_-YFP activity overshot its steady state activity. Overshooting cells were sick and also wider. Thus overshooting cells could be removed based on their width. First the single-cell width traces were smoothed with a gaussian filter and their maximum value was determined. All single cell traces with a maximum width larger than the mean width + 6 sigma of cells before the stress were removed (Figure S9 & Figure S10).

#### Growth Rate

The instantaneous growth rate as shown in Figure S11 was calculated as in (Martins et al. 2018):

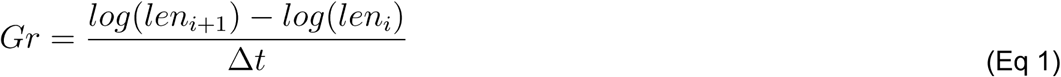

where len_i_ is the length of the cell at frame i and Δt is the time between subsequent frames taken from the image metadata. Growth rates were only calculated within one cell cycle and not over a division.

#### Calculation of surviving cells in priming experiment

The fraction of primed cells which survived the 20 µg/ml lysozyme stress treatment in Figure 2 were defined as followed:

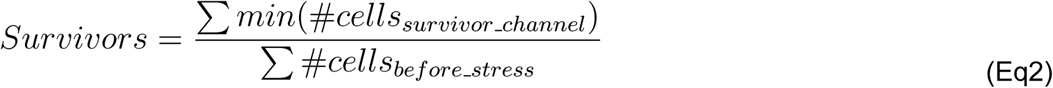

Where *Survivors* is the fraction of cells that survive the 20 µg/ml lysozyme stress treatment. *min(#cells*_*survivor_channel*_) is the smallest number of cells in a channel during the movie for channels that still have at least one cell at the end of the movie (i.e to estimate the number of surviving cells we calculated the smallest number of cells in the channel between first frame after addition of 20 µg/ml lysozyme and the end of the movie). *#cells*_*before_stress*_ is the number of cells in a channel in the last frame before addition of 20 µg/ml lysozyme stress.

#### Statistical analysis of the effects of higher P_sigV_-YFP levels before lysozyme stress application on response time

As the distributions in Figure S14 were not normal we use the non-parametric Kolmogorov–Smirnov test (ks-test) to verify the null hypothesis that cells with higher P_sigV_-YFP values before lysozyme stress application would have the same activation time distribution as cells with lower P_sigV_-YFP expression. Cells with high P_sigV_-YFP were defined as cells with a YFP fluorescence higher than the mean YFP fluorescence before the addition of lysozyme plus one standard deviation. All other cells were defined as low P_sigV_-YFP. A p-value below 0.05 was used to reject the null hypothesis. The ks-test was performed using the built in MATLAB function *kstest2*.

We found that for 0.5, 1 and 2 μg/ml lysozyme the null hypothesis was rejected. Thus cells with higher P_sigV_-YFP values before lysozyme stress application had a shorter activation time.

For 4 μg/ml lysozyme the null hypothesis was accepted. Therefore cells with higher P_sigV_-YFP values before lysozyme stress application had the same activation time as cells with lower P_sigV_-YFP values.

### RNA-seq experiment

#### RNA extraction and library preparation

10 ml of *B. subtilis* at OD 0.1-0.2 were centrifuged in 15 ml falcon tubes for 10 min at 4000 rpm. Cell pellets were resuspended in 1 ml of Qiagen RNAprotect Bacteria Reagent, and flash frozen in liquid nitrogen.

Defrosted cells were pelleted by a quick centrifugation. After removing the supernatant, cells were resuspended in 1 ml buffer RLT from Qiagen RNeasy kit. Resuspended cells were transferred in a Fastprep Lysing Matrix B tube (MP Bio) and processed in Fastprep apparatus 45 seconds at speed 6.5M/s. 700 µl of the supernatant, containing lysed cells, was transferred to a new microcentrifuge tube, to which 500 µl of absolute ethanol was added. After vortexing, the lysate was transferred to a RNeasy spin column and centrifuged 15 seconds at > 10000 rpm. RNA purification was then carried out following the instructions of the Quiagen RNeasy kit. RNA quality and integrity were assessed on the Agilent 2200 TapeStation, and RNA concentration was assessed using Qubit RNA HS assay kit. Library preparation was performed using ScriptSeq™ Complete Kit (Illumina, BB1224), for 2 µg of high integrity total RNA (RIN>8). The libraries were sequenced on a NextSeq500 using paired-end sequencing of 75 bp in length.

#### RNA-seq analysis

The raw reads were analysed using a combination of publicly available software and in-house scripts. We first assessed the quality of reads using FastQC (www.bioinformatics.babraham.ac.uk/projects/fastqc/). Reads were aligned to the Bacillus subtilis subsp. subtilis str. 168 transcriptome (http://bacteria.ensembl.org/Bacillus_subtilis_subsp_subtilis_str_168/Info/Index) using Salmon (Patro et al. 2017). Read counts for each gene were imported using the tximport R package (Soneson et al. 2015). Genes differentially expressed between the WT and *ΔsigV* mutant or in response to lysozyme treatment were identified using the DESeq2 R package (Love et al., 2014).

### Mathematical Model

We constructed a minimal mathematical model of the σ^V^ circuit that was based on the core components that we found experimentally to modulate sigV heterogeneity (Figure 3). The model consists of σ^V^, RsiV, and the σ^V^-RsiV complex. Free σ^V^ that is not bound in a complex activates the production of both σ^V^ and RsiV. Lysozyme stress was modelled as activating the cleavage of the σ^V^-RsiV complex, releasing σ^V^ while degrading RsiV. We note that although heterogeneous activation can be simulated through just a positive feedback loop, our model is the simplest model that can reproduce the perturbations of *rsiV* and *sigV* that we observe experimentally.

#### Model reactions

We used the following set of biochemical reactions to model the dynamics of σ^V^, RsiV, and σ^V^-RsiV.

##### Production

Ø -> σ^V^, RsiV

The production rate of σ^V^ and RsiV was assumed to follow a hill function, where the production rate of both σ^V^ and RsiV were identical and activated by σ^V^.

##### Dilution/Degradation

σ^V^, RsiV, σ^V^-RsiV -> Ø

We assumed a constant and identical dilution rate for all three components of the system.

##### Binding/Dissociation

σ^V^ + RsiV <-> σ^V^-RsiV

We assumed that σ^V^ and RsiV would bind to each other to form a complex and that this complex could dissociate. We also assumed the binding rate to be much higher than the dissociation rate.

##### Cleavage

σ^V^-RsiV -> σ^V^

We assumed that the σ^V^/RsiV complex would be cleaved by lysozyme, with σ^V^ getting released and RsiV degraded.

#### Model equations

The model was implemented using mass action kinetics, with a hill function representing the production rates. This generated the following system of ordinary differential equations:

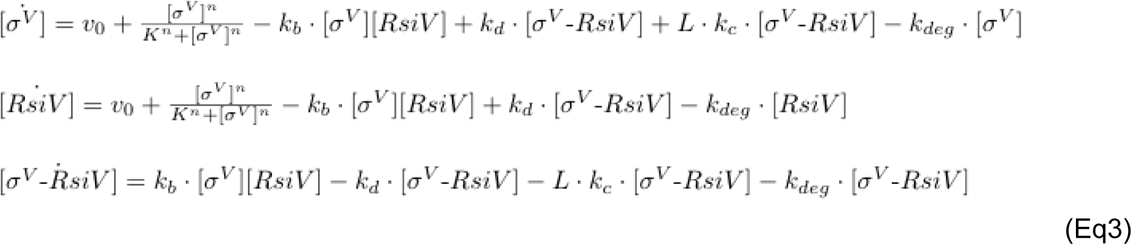

A stochastic version of the model was implemented using the chemical Langevin equations. This generated the following system of stochastic differential equations:

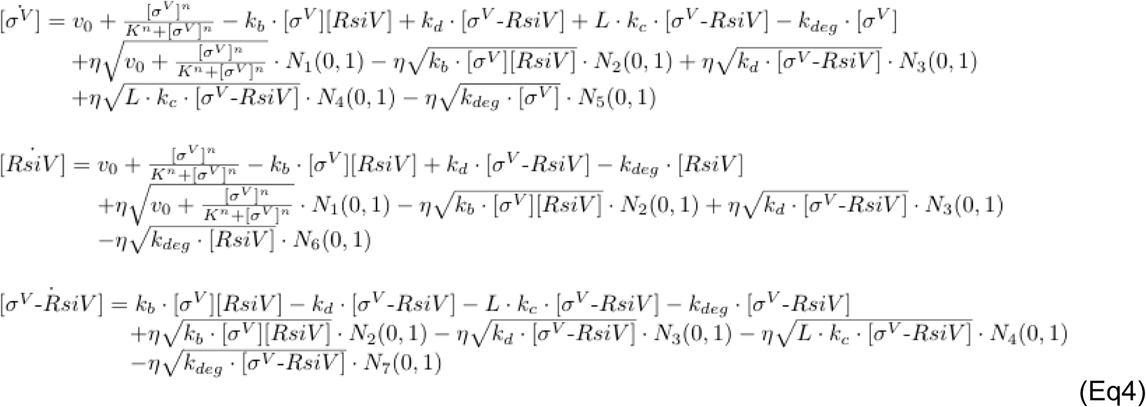

#### Model implementation

The model was implemented in the Julia programming language using the DiffEqBiological.jl modelling package. Simulations were made using the DifferentialEquations.jl package. See table S2 for more information on model parameters.

#### A note on the deterministic and stochastic regimes of the model

While the stochastic version of the bifurcation diagram (Figure S19B) only exhibits a bistable type behaviour in a finite region, we note that for these parameter values, in the deterministic bifurcation diagram bistability remains as the stress levels approach infinity (Figure S19A). Thus, a deterministic version of the model with these parameters is unable to turn on, no matter the level of lysozyme stress, as stochastic fluctuations in the expression of the *sigV* operon are required for activation. Using a modified parameter set, we verified that heterogeneous activation dynamics are still observed in the stochastic model, even when the corresponding deterministic bifurcation diagram has a finite bistability region (Figure S19).

## Code availability

Code used for simulations can be found here: https://gitlab.com/slcu/teamJL/schwall_etal_2020. Code for analysis of data reported in this study is available upon request from the corresponding author.

## Acknowledgements

We thank John Sauls and Professor Suckjoon Jun for the kind gift of a Mother Machine epoxy mould. We thank Jin Park for help with the mother machine analysis code, Michael Elowtiz for sharing plasmids, Craig Ellermeier for sharing strains, critical reading of the manuscript and helpful discussions, Katie Abley for critical reading of the manuscript and Niklas Korsbo for helpful discussions. This research was made possible by the award of a European Research Council under the European Union’s Seventh Framework Programme (FP/2007-2013)/ERC Grant Agreement 338060. The work in the Locke laboratory is further supported by a fellowship from the Gatsby Foundation (GAT3272/GLC). Torkel Loman has received funding from the European Union’s Horizon 2020 research and innovation programme under the Marie Skłodowska-Curie grant agreement No. 721456.

