## Supplementary Text for "Tunable phenotypic variability through an autoregulatory alternative sigma factor circuit"

### Supplementary Information

#### Movie Legends

All movies can be found here: https://gitlab.com/slcu/teamJL/schwall_etal_2020

**Movie S1. $\sigma^V$ is activated heterogeneously in response to lysozyme stress.** JLB130 is grown in the mother machine microfluidic device. The constitutively expressed RFP (magenta) and $P_{sigV}$-YFP(green) ranges were chosen for display. The imaging interval is 10 minutes.

**Movie S2. After a 6 h break in stress, $\sigma^V$ activation is homogenous.** JLB221 is grown in the mother machine microfluidic device. The cells experience lysozyme stress from the beginning of the movie. At time point 0 min stress is removed and then readded 6 h later. The constitutively expressed RFP (magenta) and $P_{sigV}$-YFP(green) ranges were chosen for display. The imaging interval is 10 minutes.

**Movie S3. After a 12 h break in stress, $\sigma^V$ is activated heterogeneously.** JLB221 is grown in the mother machine microfluidic device. The cells experience lysozyme stress from the beginning of the movie. At time point 0 min stress is removed and then readded 12 h later. The constitutively expressed RFP (magenta) and $P_{sigV}$-YFP(green) ranges were chosen for display. The imaging interval is 10 minutes.

<u>**Tables**</u>

**Table S1. Strain list**

| Strain Name | Genotype | Source |
|---|---|---|
| PY79 | BGSC 1A747 | BGSC |
| JP1 | PY79 ppsb::PtrpE-mCh PhleoR | (Park et al. 2018) |
| JP2 | JP1 ytvA::NeoR deletion | (Park et al. 2018) |
| JLB047 | JJB176 sacA::PsigV-yfp CmR ytvA::neoR | This work |
| JLB130 | JLB047 haG::ErmR deletion | This work |
| JLB154 | JLB130 sigV::TetR deletion | This work |
| JLB193 | JLB130 amyE::hyperspank rsiV SpecR | This work |
| JLB210 | JLB130 amyE::PsigV-sigV SpecR | This work |
| JLB211 | JLB130 amyE::PsigV-sigVrsiV SpecR | This work |
| JLB212 | JLB130 amyE::PsigV-rsiV SpecR | This work |
| JLB213 | JLB130 amyE::PsigV-mTurq SpecR | This work |
| JLB215 | JLB130 amyE::hyperspank yrhK SpecR | This work |
| JLB216 | JLB130 amyE::hyperspank sipS SpecR | This work |
| JLB217 | JLB130 amyE::hyperspank rasP SpecR | This work |
| JLB218 | JLB130 amyE::hyperspank oatA SpecR | This work |
| JLB219 | JLB130 amyE::hyperspank sigV SpecR | This work |
| JLB221 | JLB130 epsH::TetR deletion | This work |

**Table S2. Model parameters**

Parameter values for the model were set as follows.

| Parameter | Description | Value |
|---|---|---|
| $v_0$ | Leakage of the operon | 0.005 $au \cdot min^{-1}$ |
| v | Maximal operon activity | 0.1 $au \cdot min^{-1}$ |
| K | Apparent dissociation constant | 2.8 au |
| n | Hill coefficient | 4 |
| $k_D$ | Dissociation rate | 10 $min^{-1}$ |
| $k_B$ | Binding rate | 100 $au^{-1} \cdot min^{-1}$ |
| $k_C$ | Cleavage rate | 0.1 $min^{-1}$ |
| $k_{deg}$ | Dilution/degradation rate | 0.01 $min^{-1}$ |
| L | Lysozyme stress strength | variable |
| $\eta$ | Noise scaling | 0.3 |

**Table S3. Number of Repeat**

| Figure | Biological Repeats (n) | Cells (N) | Strain |
|---|---|---|---|
| 1D | 2 | 95 | JLB130 |
| 1E | 2 | 95 | JLB130 |
| 1F | 1<br>The second repeat is in Figure S 7 | 0.5 µg/ml: 45<br>1.0 µg/ml: 58<br>2.0 µg/ml: 62<br>4.0 µg/ml: 53 | JLB130 |
| 1G | 2 | 0.0 µg/ml Repeat 1: 61<br>0.0 µg/ml Repeat 2: 50<br>0.5 µg/ml Repeat 1: 45<br>0.5 µg/ml Repeat 2: 50<br>1.0 µg/ml Repeat 1: 58<br>1.0 µg/ml Repeat 2: 51<br>2.0 µg/ml Repeat 1: 62<br>2.0 µg/ml Repeat 2: 52<br>4.0 µg/ml Repeat 1: 53<br>4.0 µg/ml Repeat 2: 47 | JLB130 |
| 2B | No Pre-Stress: n=2<br>Pre-Stress: n=4 | No Priming Repeat 1: 2318<br>No Priming Repeat 2: 2209<br>Priming Repeat 1: 2237<br>Priming Repeat 2: 1811<br>Priming Repeat 3: 2027<br>Priming Repeat 4: 3773 | JLB130 |
| 2C | n=4<br>The shown data is corrected data only | Survivors Repeat 1: 46<br>Perishers Repeat 1: 77<br>Survivors Repeat 2: 30<br>Perishers Repeat 2: 49<br>Survivors Repeat 3: 120<br>Perishers Repeat 3: 122<br>Survivors Repeat 4: 54<br>Perishers Repeat 4: 66 | JLB130 |
| 3B | n=2 | Not applicable | JLB130 |
| 3C | n=2 | Not applicable | JLB154 |
| 3D | n≥3 | WT Repeat 1: 2264<br>WT Repeat 2: 3150<br>WT Repeat 3: 4480<br>sigV+ Repeat 1: 1802<br>sigV+ Repeat 2: 2386<br>sigV+ Repeat 3: 1963<br>rsiV+ Repeat 1: 2919 | WT: JLB130<br>SigV: JLB219<br>RsiV: JLB193<br>OatA: JLB218<br>YrhK: JLB215<br>SipS: JLB216<br>RasP: JLB217 |

| | | | |
|---|---|---|---|
| | | rsiV+ Repeat 2: 2098<br>rsiV+ Repeat 3: 2397<br>oatA+ Repeat 1: 3025<br>oatA+ Repeat 2: 2147<br>oatA+ Repeat 3: 2449<br>yrhK+ Repeat 1: 2130<br>yrhK+ Repeat 2: 2585<br>yrhK+ Repeat 3: 3071<br>sipS+ Repeat 1: 914<br>sipS+ Repeat 2: 2999<br>sipS+ Repeat 3: 4081<br>rasP+ Repeat 1: 2741<br>rasP+ Repeat 2: 3194<br>rasP+ Repeat 3: 2970 | |
| 3E | n=3 | WT 0 Lyso & 1 IPTG Repeat 1: 2820<br>WT 0 Lyso & 1 IPTG Repeat 2: 2529<br>WT 0 Lyso & 1 IPTG Repeat 3: 2012<br>WT 1 Lyso & 1 IPTG Repeat 1: 3938<br>WT 1 Lyso & 1 IPTG Repeat 2: 2677<br>WT 1 Lyso & 1 IPTG Repeat 3: 2324<br>oatA+ 1 Lyso & 1 IPTG Repeat 1: 2147<br>oatA+ 1 Lyso & 1 IPTG Repeat 2: 2449<br>oatA+ 1 Lyso & 1 IPTG Repeat 3: 4200<br>oatA+ 20 Lyso & 1 IPTG Repeat 1: 2730<br>oatA+ 20 Lyso & 1 IPTG Repeat 2: 2501<br>oatA+ 20 Lyso & 1 IPTG Repeat 3: 2906 | WT: JLB130<br>OatA: JLB218 |
| 4A | Not applicable | 100 | Not applicable |
| 4B | Not applicable | 999 | Not applicable |
| 4C | Not applicable | 999 | Not applicable |
| 5B | 1000 bootstraps | 999 | Not applicable |
| 5C | 3 | WT Repeat 1: 499<br>WT Repeat 2: 624<br>WT Repeat 3: 365<br>2x sigV Repeat 1: 454<br>2x sigV Repeat 2: 443<br>2x sigV Repeat 3: 400<br>2x rsiV Repeat 1: 510<br>2x rsiV Repeat 2: 674<br>2x rsiV Repeat 3: 502<br>2x sigVrsiV Repeat 1: 434<br>2x sigVrsiV Repeat 2: 475<br>2x sigVrsiV Repeat 3: 342 | WT: JLB130<br>2xsigV: JLB210<br>2xrsiV: JLB212<br>2xsigVrsiV: JLB211 |

| | | | |
|---|---|---|---|
| 6A | Not applicable | 100 | Not applicable |
| 6B | Not applicable | 100 | Not applicable |
| 6C | Not applicable | 300 au break: 500<br>450 au break: 500<br>600 au break: 500 | Not applicable |
| 6D | 1 the repeat is in Figure S24 | 48 | JLB221 |
| 6E | 1 the repeat is in Figure S24 | 54 | JLB221 |
| 6F | 1 the repeat is in Figure S24 | 6h no stress: 48<br>12h no stress: 54<br>Control: 56 | JLB221 |
| S1 | 2 all repeats are shown in the figure | A) 0.5 µg/ml Repeat 1: 45<br>B) 0.5 µg/ml Repeat 2: 50<br>C) 1.0 µg/ml Repeat 1: 58<br>D) 1.0 µg/ml Repeat 2: 51<br>E) 2.0 µg/ml Repeat 1: 62<br>F) 2.0 µg/ml Repeat 2: 52<br>G) 4.0 µg/ml Repeat 1: 53<br>H) 4.0 µg/ml Repeat 2: 47 | JLB130 |
| S2 | 2 all repeats are shown in the figure | A) 0.5 µg/ml Repeat 1: 45<br>B) 0.5 µg/ml Repeat 2: 50<br>C) 1.0 µg/ml Repeat 1: 58<br>D) 1.0 µg/ml Repeat 2: 51<br>E) 2.0 µg/ml Repeat 1: 62<br>F) 2.0 µg/ml Repeat 2: 52<br>G) 4.0 µg/ml Repeat 1: 53<br>H) 4.0 µg/ml Repeat 2: 47 | JLB130 |
| S3 | 2 | A) 0.5 µg/ml: 95<br>B) 1.0 µg/ml: 109<br>C) 2.0 µg/ml: 114<br>D) 4.0 µg/ml: 100 | JLB130 |
| S4 | Movie data (red lines): 2<br>Snap data (black lines): 8 | Movie Data Repeat 1: 766<br>Movie Data Repeat 2: 744<br>Snap Data Repeat 1: 3144<br>Snap Data Repeat 2: 3443<br>Snap Data Repeat 3: 2901<br>Snap Data Repeat 4: 2667<br>Snap Data Repeat 5: 3736<br>Snap Data Repeat 6: 3090 | JLB130 |

| | | | |
|---|---|---|---|
| | | Snap Data Repeat 7: 2687<br>Snap Data Repeat 8: 4239 | |
| S5 | 3 | Cell at the end of the movie:<br>A) 0.0 µg/ml: 1003<br>B) 0.5 µg/ml: 626<br>C) 1.0 µg/ml: 573<br>D) 4.0 µg/ml: 825 | JLB47 |
| S6 | 3 | 3238 | JLB213 |
| S7 | 1 the second repeat is in figure 1E | 0.5 µg/ml: 50<br>1.0 µg/ml: 51<br>2.0 µg/ml: 52<br>4.0 µg/ml: 47 | JLB130 |
| S8 | 2 all repeats are shown in the figure | A) 0.5 µg/ml Repeat 1: 45<br>B) 0.5 µg/ml Repeat 2: 50<br>C) 1.0 µg/ml Repeat 1: 58<br>D) 1.0 µg/ml Repeat 2: 51<br>E) 2.0 µg/ml Repeat 1: 61<br>F) 2.0 µg/ml Repeat 2: 51<br>G) 4.0 µg/ml Repeat 1: 49<br>H) 4.0 µg/ml Repeat 2: 37 | JLB130 |
| S9 | 2:<br>Two red lines:<br>All Cells<br>Two blue lines:<br>Overshooters removed | Red lines:<br>Repeat 1: 53<br>Repeat 2: 47<br><br>Blue lines:<br>Repeat 1: 49<br>Repeat 2: 37 | JLB130 |
| S10 | 2 | A) 0.5 µg/ml: 72<br>B) 1.0 µg/ml: 94<br>C) 2.0 µg/ml: 72<br>D) 4.0 µg/ml: 53 | JLB130 |
| S11 | 2 | A) 0.5 µg/ml: 95<br>B) 1.0 µg/ml: 109<br>C) 2.0 µg/ml: 112<br>D) 4.0 µg/ml: 86 | JLB130 |
| S12 | 2 | YFP & RFP Repeat 1: 61<br>YFP & RFP Repeat 2: 50 | JLB130 |
| S13 | 2 | A) 0.5 µg/ml: 95<br>B) 1.0 µg/ml: 109<br>C) 2.0 µg/ml: 112<br>D) 4.0 µg/ml: 86 | JLB130 |

| | | | |
|---|---|---|---|
| S14 | 2 | A) 0.5 µg/ml:<br>Red: 11; Blue: 84<br>B) 1.0 µg/ml:<br>Red: 11; Blue: 98<br>C) 2.0 µg/ml:<br>Red: 15; Blue: 97<br>D) 4.0 µg/ml: 86<br>Red: 12; Blue: 74 | JLB130 |
| S15 | A) n=2<br>B) n=4 | A) N=113<br>B) Survivors: 250<br>Perishers: 326 | JLB130 |
| S16 | WT 0 Lyso & 0 IPTG n: 8<br>WT 1 Lyso & 0 IPTG n: 8<br>WT 0 Lyso & 1 IPTG n: 8<br>WT 1 Lyso & 1 IPTG n: 8<br>rsiV+ 0 Lyso & 0 IPTG n: 3<br>rsiV+ 1 Lyso & 0 IPTG n: 3<br>rsiV+ 0 Lyso & 1 IPTG n: 3<br>rsiV+ 1 Lyso & 1 IPTG n: 3<br>yrhK+ 0 Lyso & 0 IPTG n: 3<br>yrhK+ 1 Lyso & 0 IPTG n: 3<br>yrhK+ 0 Lyso & 1 IPTG n: 3<br>yrhK+ 1 Lyso & 1 IPTG n: 3<br>sipS+ 0 Lyso & 0 IPTG n: 3<br>sipS+ 1 Lyso & 0 IPTG n: 3<br>sipS+ 0 Lyso & 1 IPTG n: 3<br>sipS+ 1 Lyso & 1 IPTG n: 3<br>rasP+ 0 Lyso & 0 IPTG n: 3<br>rasP+ 1 Lyso & 0 IPTG n: 3<br>rasP+ 0 Lyso & 1 IPTG n: 3<br>rasP+ 1 Lyso & 1 IPTG n: 3<br>oatA+ 0 Lyso & 0 IPTG n: 3<br>oatA+ 1 Lyso & 0 IPTG n: 3<br>oatA+ 0 Lyso & 1 IPTG n: 3<br>oatA+ 1 Lyso & 1 IPTG n: 3<br>sigV+ 0 Lyso & 0 IPTG n: 4<br>sigV+ 1 Lyso & 0 IPTG n: 4<br>sigV+ 0 Lyso & 1 IPTG n: 4<br>sigV+ 1 Lyso & 1 IPTG n: 4 | WT 0 Lyso & 0 IPTG Repeat 1: 3019<br>WT 0 Lyso & 0 IPTG Repeat 2: 2720<br>WT 0 Lyso & 0 IPTG Repeat 3: 2721<br>WT 0 Lyso & 0 IPTG Repeat 4: 3690<br>WT 0 Lyso & 0 IPTG Repeat 5: 2142<br>WT 0 Lyso & 0 IPTG Repeat 6: 4295<br>WT 0 Lyso & 0 IPTG Repeat 7: 2553<br>WT 0 Lyso & 0 IPTG Repeat 8: 4295<br>WT 1 Lyso & 0 IPTG Repeat 1: 3072<br>WT 1 Lyso & 0 IPTG Repeat 2: 3012<br>WT 1 Lyso & 0 IPTG Repeat 3: 2752<br>WT 1 Lyso & 0 IPTG Repeat 4: 2543<br>WT 1 Lyso & 0 IPTG Repeat 5: 3710<br>WT 1 Lyso & 0 IPTG Repeat 6: 3077<br>WT 1 Lyso & 0 IPTG Repeat 7: 2647<br>WT 1 Lyso & 0 IPTG Repeat 8: 4182<br>WT 0 Lyso & 1 IPTG Repeat 1: 3186<br>WT 0 Lyso & 1 IPTG Repeat 2: 2820<br>WT 0 Lyso & 1 IPTG Repeat 3: 2139<br>WT 0 Lyso & 1 IPTG Repeat 4: 3325<br>WT 0 Lyso & 1 IPTG Repeat 5: 2529<br>WT 0 Lyso & 1 IPTG Repeat 6: 2898<br>WT 0 Lyso & 1 IPTG Repeat 7: 2012<br>WT 0 Lyso & 1 IPTG Repeat 8: 2839<br>WT 1 Lyso & 1 IPTG Repeat 1: 3150<br>WT 1 Lyso & 1 IPTG Repeat 2: 3938<br>WT 1 Lyso & 1 IPTG Repeat 3: 2264<br>WT 1 Lyso & 1 IPTG Repeat 4: 4493<br>WT 1 Lyso & 1 IPTG Repeat 5: 2677<br>WT 1 Lyso & 1 IPTG Repeat 6: 2832<br>WT 1 Lyso & 1 IPTG Repeat 7: 2324<br>WT 1 Lyso & 1 IPTG Repeat 8: 4480<br>rsiV+ 0 Lyso & 0 IPTG Repeat 1: 3264<br>rsiV+ 0 Lyso & 0 IPTG Repeat 2: 2319<br>rsiV+ 0 Lyso & 0 IPTG Repeat 3: 1399<br>rsiV+ 1 Lyso & 0 IPTG Repeat 1: 2734<br>rsiV+ 1 Lyso & 0 IPTG Repeat 2: 3317<br>rsiV+ 1 Lyso & 0 IPTG Repeat 3: 2148 | WT: JLB130<br>SigV: JLB219<br>RsiV: JLB193<br>OatA: JLB218<br>YrhK: JLB215<br>SipS: JLB216<br>RasP: JLB217 |

| | | |
|---|---|---|
| | | rsiV+ 0 Lyso & 1 IPTG Repeat 1: 2300<br>rsiV+ 0 Lyso & 1 IPTG Repeat 2: 2612<br>rsiV+ 0 Lyso & 1 IPTG Repeat 3: 1856<br>rsiV+ 1 Lyso & 1 IPTG Repeat 1: 2919<br>rsiV+ 1 Lyso & 1 IPTG Repeat 2: 2098<br>rsiV+ 1 Lyso & 1 IPTG Repeat 3: 2397<br>yrhK+ 0 Lyso & 0 IPTG Repeat 1: 3088<br>yrhK+ 0 Lyso & 0 IPTG Repeat 2: 3295<br>yrhK+ 0 Lyso & 0 IPTG Repeat 3: 3282<br>yrhK+ 1 Lyso & 0 IPTG Repeat 1: 2116<br>yrhK+ 1 Lyso & 0 IPTG Repeat 2: 3285<br>yrhK+ 1 Lyso & 0 IPTG Repeat 3: 2936<br>yrhK+ 0 Lyso & 1 IPTG Repeat 1: 2107<br>yrhK+ 0 Lyso & 1 IPTG Repeat 2: 936<br>yrhK+ 0 Lyso & 1 IPTG Repeat 3: 2528<br>yrhK+ 1 Lyso & 1 IPTG Repeat 1: 2130<br>yrhK+ 1 Lyso & 1 IPTG Repeat 2: 2585<br>yrhK+ 1 Lyso & 1 IPTG Repeat 3: 3071<br>sipS+ 0 Lyso & 0 IPTG Repeat 1: 2221<br>sipS+ 0 Lyso & 0 IPTG Repeat 2: 3401<br>sipS+ 0 Lyso & 0 IPTG Repeat 3: 3940<br>sipS+ 1 Lyso & 0 IPTG Repeat 1: 2478<br>sipS+ 1 Lyso & 0 IPTG Repeat 2: 2362<br>sipS+ 1 Lyso & 0 IPTG Repeat 3: 3683<br>sipS+ 0 Lyso & 1 IPTG Repeat 1: 2683<br>sipS+ 0 Lyso & 1 IPTG Repeat 2: 2837<br>sipS+ 0 Lyso & 1 IPTG Repeat 3: 3008<br>sipS+ 1 Lyso & 1 IPTG Repeat 1: 914<br>sipS+ 1 Lyso & 1 IPTG Repeat 2: 2999<br>sipS+ 1 Lyso & 1 IPTG Repeat 3: 4081<br>rasP+ 0 Lyso & 0 IPTG Repeat 1: 3499<br>rasP+ 0 Lyso & 0 IPTG Repeat 2: 2561<br>rasP+ 0 Lyso & 0 IPTG Repeat 3: 3482<br>rasP+ 1 Lyso & 0 IPTG Repeat 1: 2645<br>rasP+ 1 Lyso & 0 IPTG Repeat 2: 1607<br>rasP+ 1 Lyso & 0 IPTG Repeat 3: 3626<br>rasP+ 0 Lyso & 1 IPTG Repeat 1: 2133<br>rasP+ 0 Lyso & 1 IPTG Repeat 2: 2081<br>rasP+ 0 Lyso & 1 IPTG Repeat 3: 3466<br>rasP+ 1 Lyso & 1 IPTG Repeat 1: 2741<br>rasP+ 1 Lyso & 1 IPTG Repeat 2: 3194<br>rasP+ 1 Lyso & 1 IPTG Repeat 3: 2970<br>oatA+ 0 Lyso & 0 IPTG Repeat 1: 3763<br>oatA+ 0 Lyso & 0 IPTG Repeat 2: 3170<br>oatA+ 0 Lyso & 0 IPTG Repeat 3: 3342<br>oatA+ 1 Lyso & 0 IPTG Repeat 1: 2783<br>oatA+ 1 Lyso & 0 IPTG Repeat 2: 3216<br>oatA+ 1 Lyso & 0 IPTG Repeat 3: 4297<br>oatA+ 0 Lyso & 1 IPTG Repeat 1: 3898<br>oatA+ 0 Lyso & 1 IPTG Repeat 2: 2850 |

| | | | |
|---|---|---|---|
| | | oatA+ 0 Lyso & 1 IPTG Repeat 3: 3743<br>oatA+ 1 Lyso & 1 IPTG Repeat 1: 2147<br>oatA+ 1 Lyso & 1 IPTG Repeat 2: 2449<br>oatA+ 1 Lyso & 1 IPTG Repeat 3: 4200<br>sigV+ 0 Lyso & 0 IPTG Repeat 1: 2742<br>sigV+ 0 Lyso & 0 IPTG Repeat 2: 3401<br>sigV+ 0 Lyso & 0 IPTG Repeat 3: 4468<br>sigV+ 0 Lyso & 0 IPTG Repeat 4: 4637<br>sigV+ 1 Lyso & 0 IPTG Repeat 1: 3362<br>sigV+ 1 Lyso & 0 IPTG Repeat 2: 3510<br>sigV+ 1 Lyso & 0 IPTG Repeat 3: 3515<br>sigV+ 1 Lyso & 0 IPTG Repeat 4: 2947<br>sigV+ 0 Lyso & 1 IPTG Repeat 1: 906<br>sigV+ 0 Lyso & 1 IPTG Repeat 2: 2122<br>sigV+ 0 Lyso & 1 IPTG Repeat 3: 1647<br>sigV+ 0 Lyso & 1 IPTG Repeat 4: 1721<br>sigV+ 1 Lyso & 1 IPTG Repeat 1: 1963<br>sigV+ 1 Lyso & 1 IPTG Repeat 2: 2386<br>sigV+ 1 Lyso & 1 IPTG Repeat 3: 1802<br>sigV+ 1 Lyso & 1 IPTG Repeat 4: 1669 | |
| S17 | Not applicable | 100 | Not applicable |
| S18 | A) Not applicable<br>B) Not applicable | A) Not applicable<br>B) N= 100 per grid point; 80x250 grid points | Not applicable |
| S19 | Not applicable | 100 | Not applicable |
| S20 | WT 0 µg/ml n: 3<br>WT 1 µg/ml n: 3<br>2x sigV 0 µg/ml n: 3<br>2x sigV 1 µg/ml n: 3<br>2x rsiV 0 µg/ml n: 3<br>2x rsiV 1 µg/ml n: 3<br>2x sigV-rsiV 0 µg/ml n: 3<br>2x sigV-rsiV 1 µg/ml n: 3 | WT 0 Lyso Repeat 1: 483<br>WT 0 Lyso Repeat 2: 673<br>WT 0 Lyso Repeat 3: 439<br>WT 1 Lyso Repeat 1: 499<br>WT 1 Lyso Repeat 2: 624<br>WT 1 Lyso Repeat 3: 365<br>2xsigV 0 Lyso Repeat 1: 495<br>2xsigV 0 Lyso Repeat 2: 353<br>2xsigV 0 Lyso Repeat 3: 280<br>2xsigV 1 Lyso Repeat 1: 454<br>2xsigV 1 Lyso Repeat 2: 443<br>2xsigV 1 Lyso Repeat 3: 400<br>2xrsiV 0 Lyso Repeat 1: 446<br>2xrsiV 0 Lyso Repeat 2: 353<br>2xrsiV 0 Lyso Repeat 3: 327<br>2xrsiV 1 Lyso Repeat 1: 510<br>2xrsiV 1 Lyso Repeat 2: 674<br>2xrsiV 1 Lyso Repeat 3: 502<br>2xsigVrsiV 0 Lyso Repeat 1: 402<br>2xsigVrsiV 0 Lyso Repeat 2: 550<br>2xsigVrsiV 0 Lyso Repeat 3: 431<br>2xsigVrsiV 1 Lyso Repeat 1: 434 | WT: JLB130<br>2x sigV:<br>JLB210<br>2x risV:<br>JLB212<br>2x sigV-rsiV:<br>JLB211 |

| | | | |
|---|---|---|---|
| | | 2xsigVrsiV 1 Lyso Repeat 2: 475<br>2xsigVrsiV 1 Lyso Repeat 3: 342 | |
| S21 | Not applicable | 100 for all subplots | Not applicable |
| S22 | 3 | WT 1L Repeat 1: 841<br>WT 1L Repeat 2: 1141<br>WT 1L Repeat 3: 961<br>ΔepsH 1L Repeat 1: 1219<br>ΔepsH 1L Repeat 2: 1617<br>ΔepsH 1L Repeat 3: 1104 | WT: JLB130<br>ΔepsH:<br>JLB221 |
| S23 | A-E) 2 | A-E)<br>WT Repeat 1: 50<br>ΔepsH Repeat 1: 50<br>WT Repeat 2: 53<br>ΔepsH Repeat 2: 56 | WT: JLB130<br>ΔepsH:<br>JLB221 |
| S24 | 1<br>The other repat is in figure 6 of the main text | 6h no stress: 48<br>12h no stress: 50<br>Control: 50 | JLB221 |
| S25 | 1 | Memory 6h: 20<br>Memory 12h: 26 | JLB130 |

#### SUPPLEMENTARY FIGURES

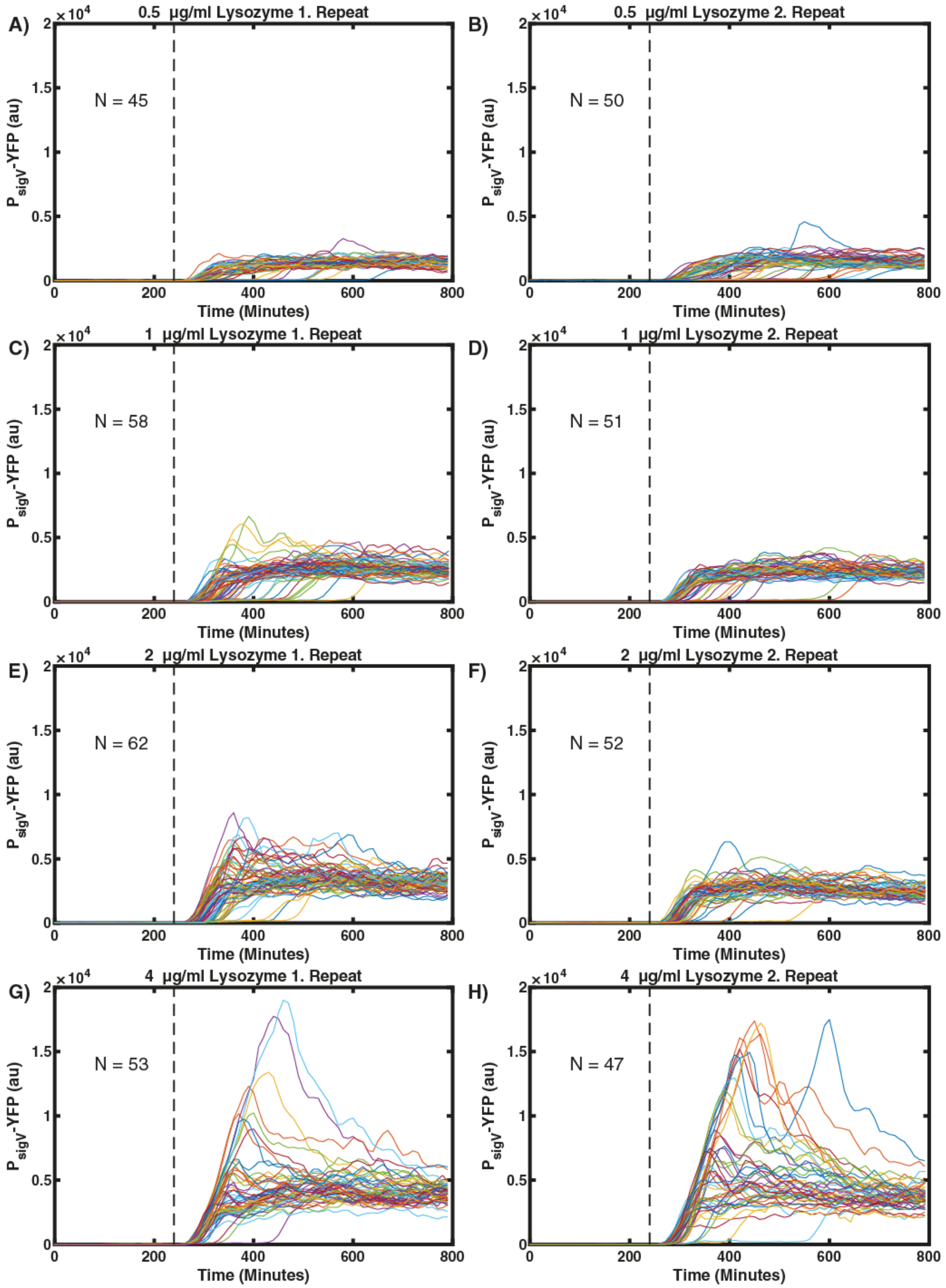

**Figure S1. Expression dynamics of $P_{sigV}$-YFP reporter under lysozyme stress.** In each subpanel a line corresponds to a single-cell trace of one mother cell in the mother machine

(N=~50). The stress was added after 240 min (dashed line). (A&B) Single cell traces in response to 0.5 µg/ml lysozyme. (C&D) Single cell traces in response to 1 µg/ml lysozyme. (E&F) Single cell traces in response to 2 µg/ml lysozyme. (G&H) Single cell traces in response to 4 µg/ml lysozyme. The shown data corresponds to the data shown in Figure 1 and Figure S2.

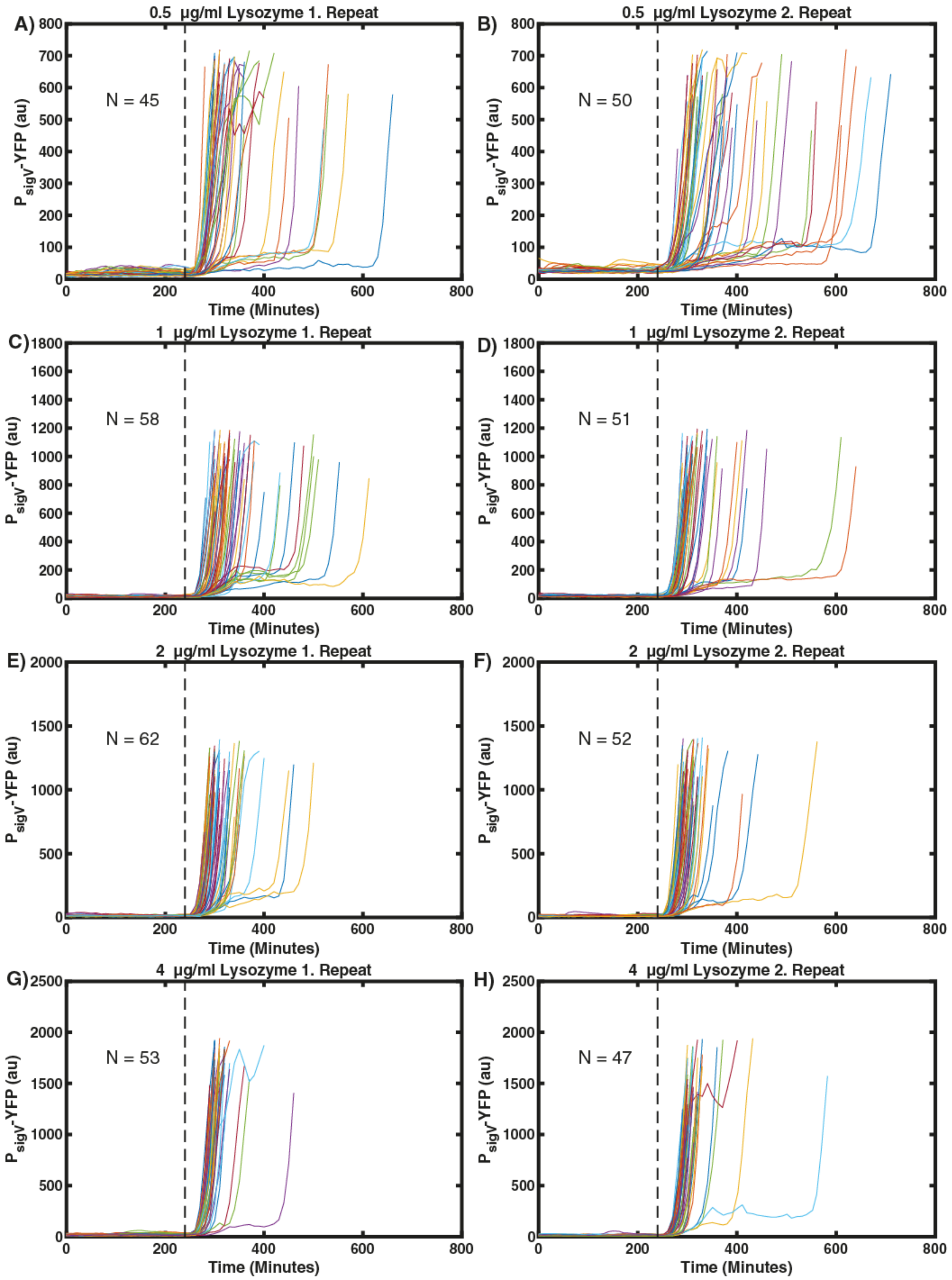

**Figure S2. With increasing stress levels the heterogeneity in $\sigma^V$ activation times are reduced.** In each subpanel a line corresponds to a single-cell trace of one mother cell in the mother machine (N=~50). The stress was added after 240 min (dashed line). Once $P_{sigV}$-YFP passed its half maximum the traces end. (A&B) Single cell traces in response to 0.5 μg/ml lysozyme. (C&D) Single cell traces in response to 1 μg/ml lysozyme. (E&F) Single cell traces in response to 2 μg/ml lysozyme. (G&H) Single cell traces in response to 4 μg/ml lysozyme. The data is the same as in Figure S1.

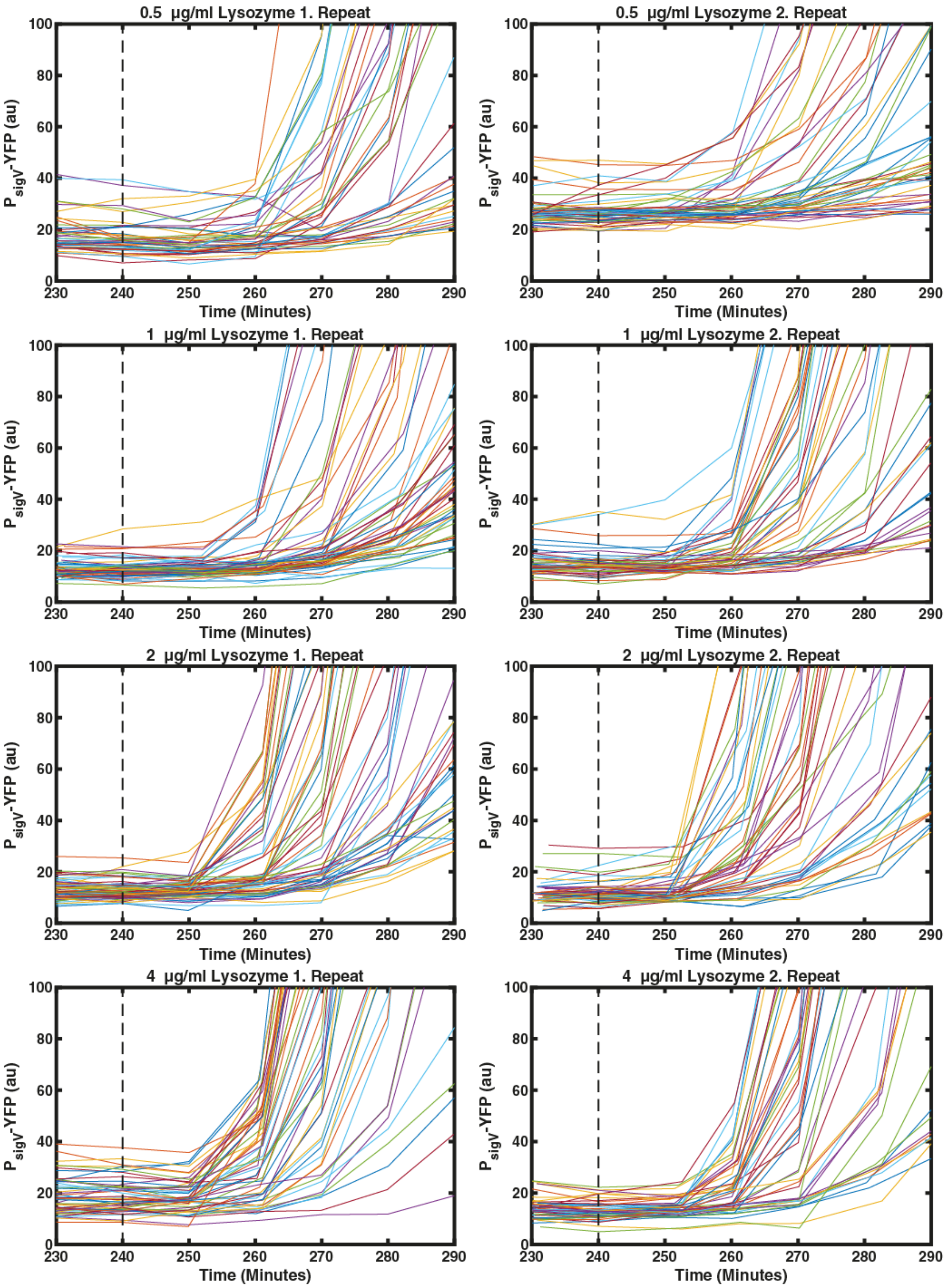

**Figure S3 A fraction of cells respond rapidly to lysozyme.** The first cells begin to respond to the lysozyme stress with raised $P_{sigV}$-YFP levels within 2 frames (20 minutes). In each subpanel a line corresponds to a single-cell trace of one mother cell in the mother machine

(N=~50). The stress was added after 240 min (dashed line). (A&B) Single cell traces in response to 0.5 µg/ml lysozyme. (C&D) Single cell traces in response to 1 µg/ml lysozyme. (E&F) Single cell traces in response to 2 µg/ml lysozyme. (G&H) Single cell traces in response to 4 µg/ml lysozyme. The shown data is the same as in Figure S1.

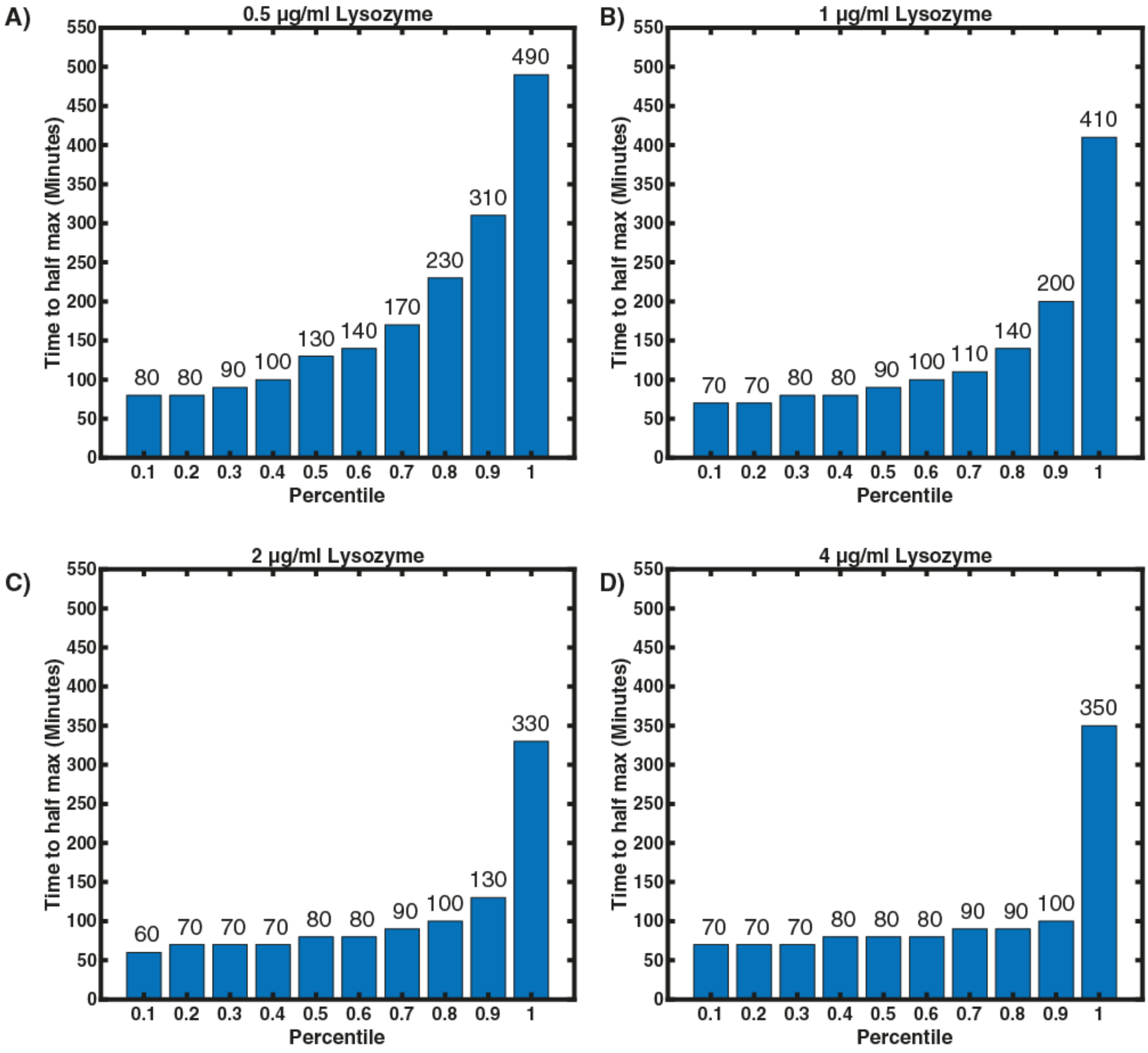

**Figure S4. With increasing stress levels it takes less time for all cells to activate the $\sigma^V$ pathway.** Each bar represents the maximum value of each $10^{th}$ percentile of activation times. The activation time was calculated as the time between switching from SMM to lysozyme and the time point when then $P_{sigV}$-YFP passed its half maximum. The number above the bar indicates the maximum activation for the corresponding 10th percentile. The subfigures show the data for N number of cells from n biological repeats. (A) $\sigma^V$ activation times in response to 0.5 μg/ml lysozyme (N=95, n=2). (B) $\sigma^V$ activation times in response to1 μg/ml lysozyme (N=109, n=2). (C) $\sigma^V$ activation times in response to 2 μg/ml lysozyme (N=114, n=2). (D) $\sigma^V$ activation times in response to 4 μg/ml lysozyme (N=100, n=2).

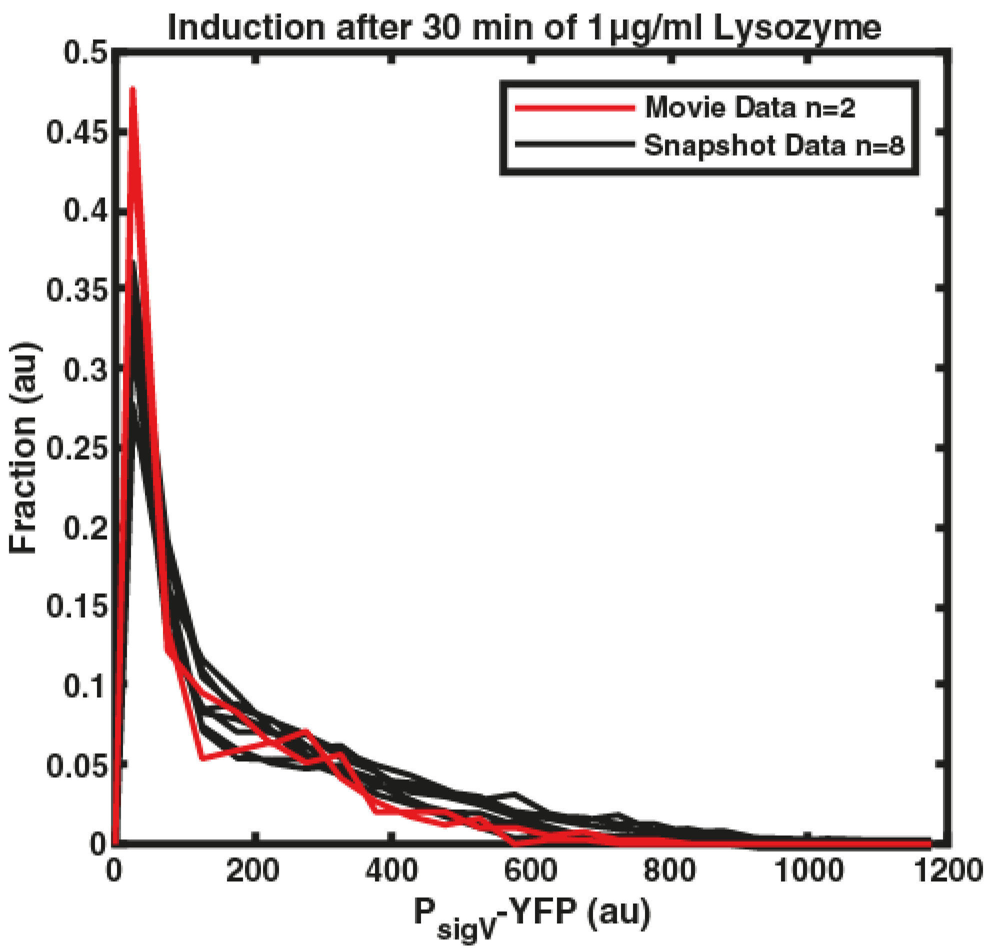

**Figure S5. Cells grown in liquid culture and the mother machine both display long tailed $\sigma^V$ activation distribution.** The black lines are the histograms of $P_{sigV}$-YFP fluorescence of cells grown in liquid culture (n=8 biological repeats with N>2600 cells each) and imaged with snapshots after being exposed to 1 µg/ml lysozyme for 30 min. The red lines are the histograms of $P_{sigV}$-YFP fluorescence of cells grown in the mother machine after being exposed to 1 µg/ml lysozyme for 30 min (n=2 with N>50 cells each). For more details on the number of cells for each biological repeat see Table S1.

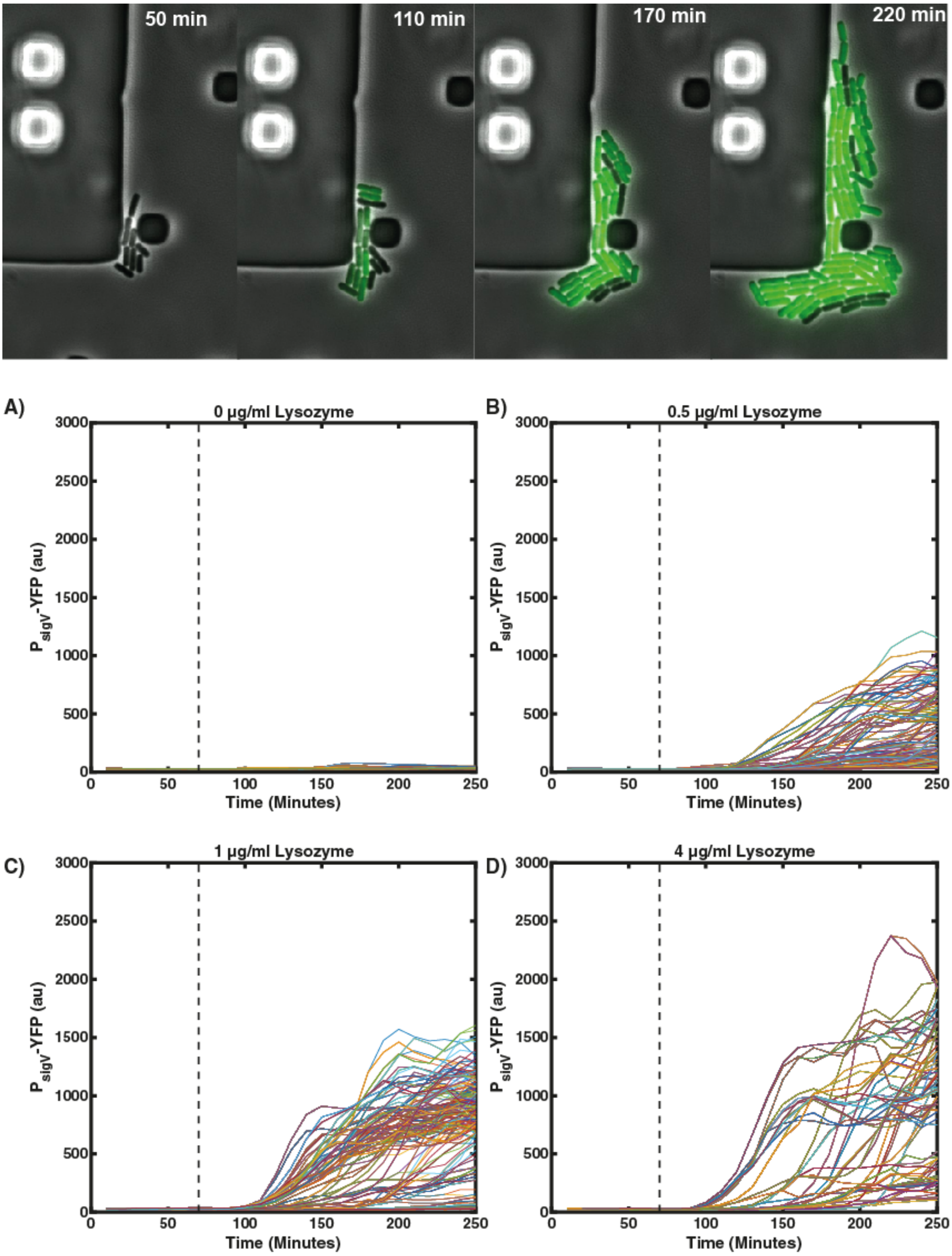

**Figure S6. The observed heterogeneity in $\sigma^V$ activation is not due to the geometry of the mother machine.** Cells grown in CellAsic bacteria chips also showed a heterogeneous activation of σ$^V$ in response to lysozyme. In each subpanel stress was added after 60 min (dashed black line) and a line corresponds to a single-cell trace. Top panel: micrographs of cells grown in the CellAsic. (A) Single cell traces in response to 0 μg/ml lysozyme. (B) Single cell traces in response to 0.5 μg/ml lysozyme. (C) Single cell traces in response to 1 μg/ml lysozyme. (D) Single cell traces in response to 4 μg/ml lysozyme. The plotted data is from three biological repeats.

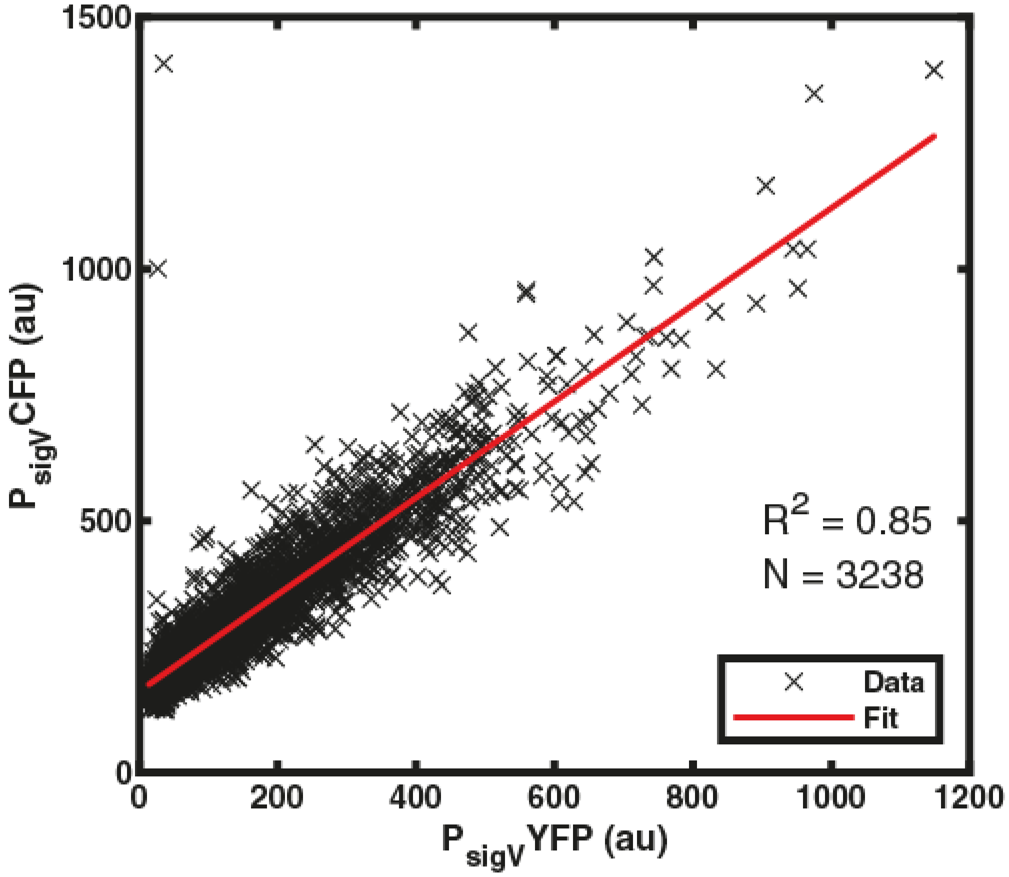

**Figure S7. The observed heterogeneity in P$_{sigV}$-YFP activation in response to lysozyme reflects a heterogeneous σ$^V$ activation and not intrinsic variability of the P$_{sigV}$-YFP promoter.** JLB213 (sacA::PsigV-yfp CmR amyE::PsigV_mTurq SpecR) was exposed to 1 μg/ml lysozyme for 30 min and imaged with snapshots. Each data point corresponds to a single cell. Data from three biological repeats are plotted together (N=3238).

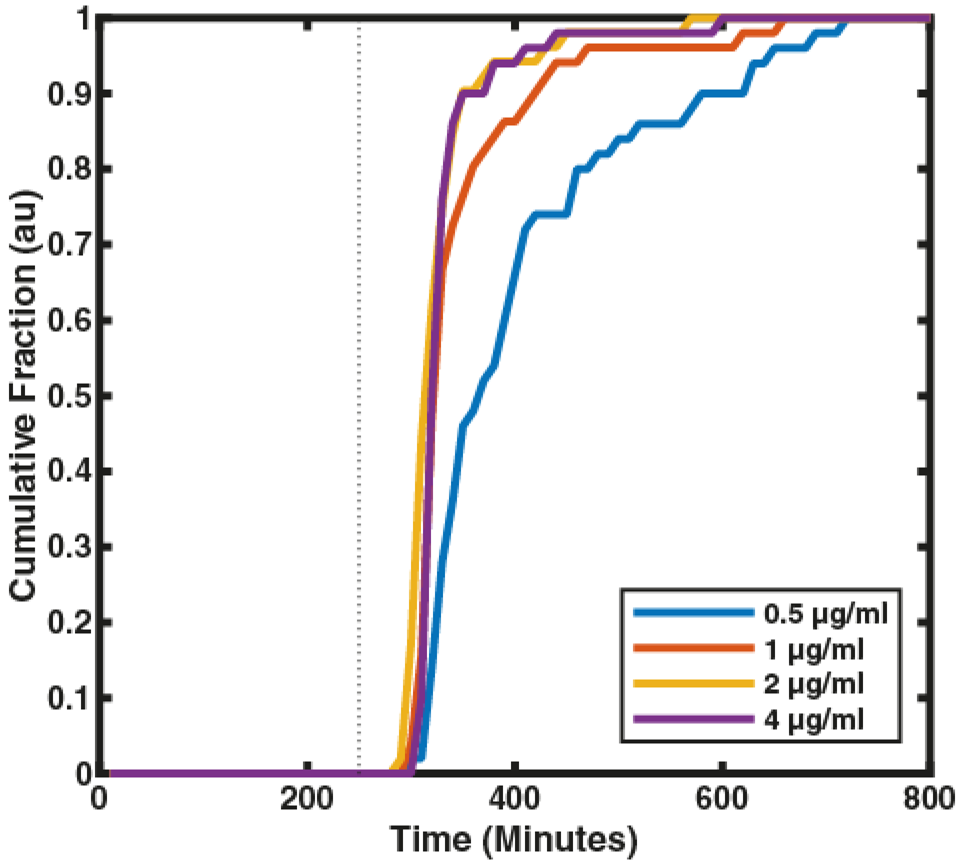

**Figure S8. With increasing stress levels the heterogeneity in $\sigma^V$ activation disappears.** The observed heterogeneity is reduced with increasing stress levels. Each line represents the time dependent fraction of single cells to activate $\sigma^V$ in response to lysozyme for N=~50 cells from one day. Data is a repeat experiment of Figure 1F.

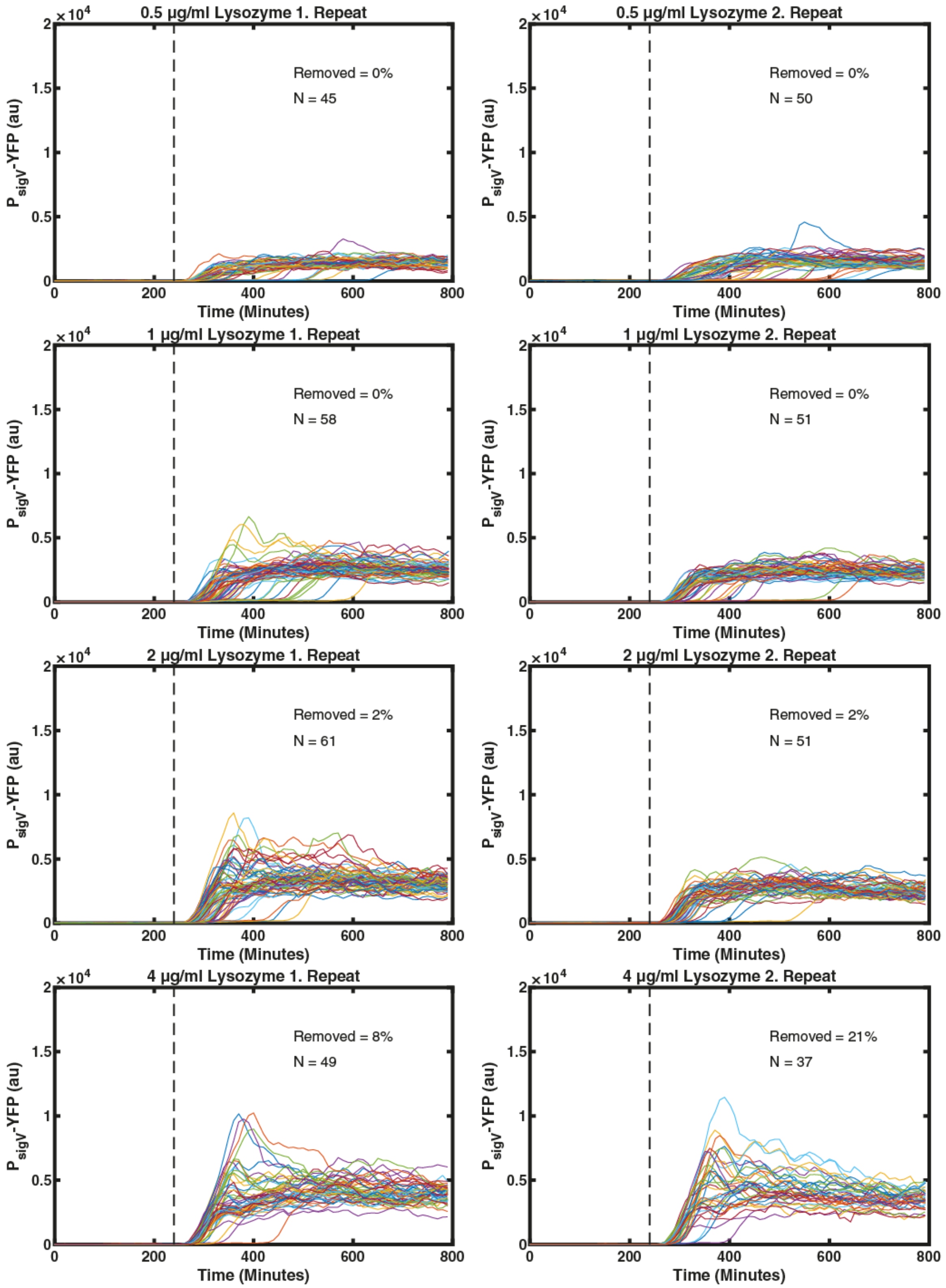

**Figure S9. Removing all wide cells from analysis removes most cells which overshoot their $\sigma^V$ activation.** Figure is the same as figure S1, but cells that are wider than unstressed cells are removed. In each subpanel a line corresponds to a single-cell trace of one mother cell in the mother machine. The stress was added after 240 min. (A&B) Single cell traces in response to 0.5 µg/ml lysozyme. (C&D) Single cell traces in response to 1 µg/ml lysozyme. (E&F) Single cell traces in response to 2 µg/ml lysozyme. (G&H) Single cell traces in response to 4 µg/ml lysozyme.

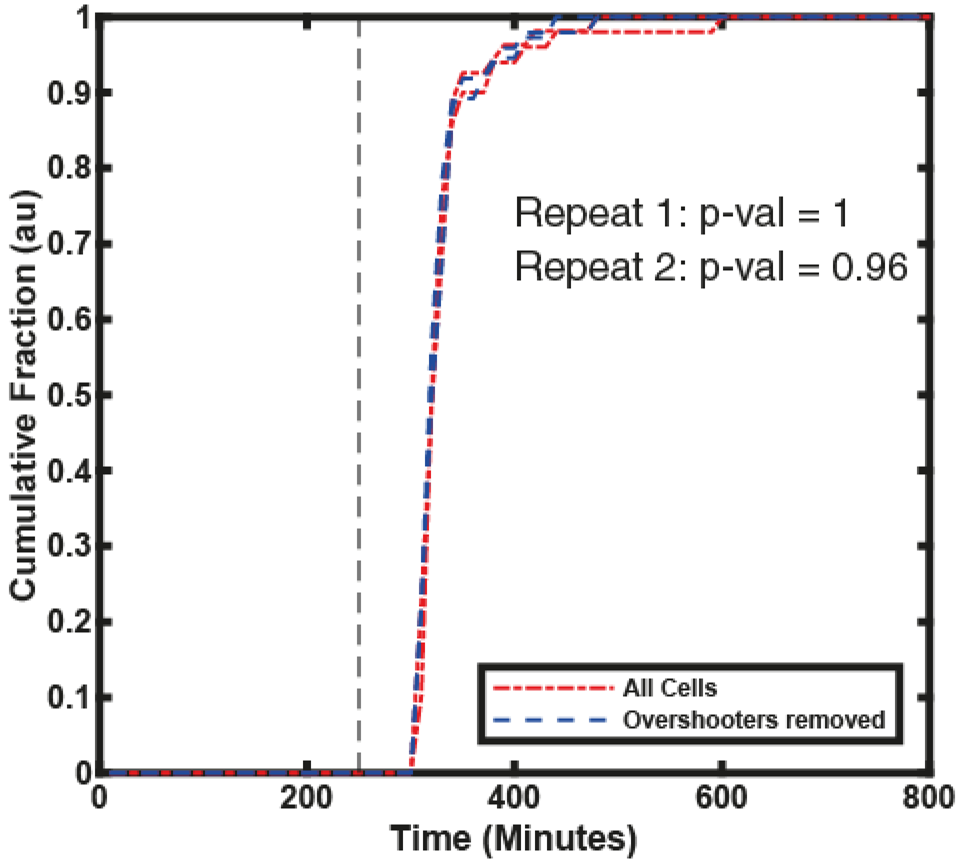

**Figure S10 Removing wide cells does not affect the observed heterogeneity.**

The cumulative fraction of cells with $P_{sigV}$-YFP values higher than the half maximum of their final values for 4 µg/ml lysozyme did not change when removing wide cells. The null hypothesis that the activation times of all cells and the activation times of cells without wide cells would have the same activation time distribution was not rejected using a Kolmogorov–Smirnov test. Only 4 µg/ml lysozyme (n=2) is shown as wide cells were not observed at other concentrations.

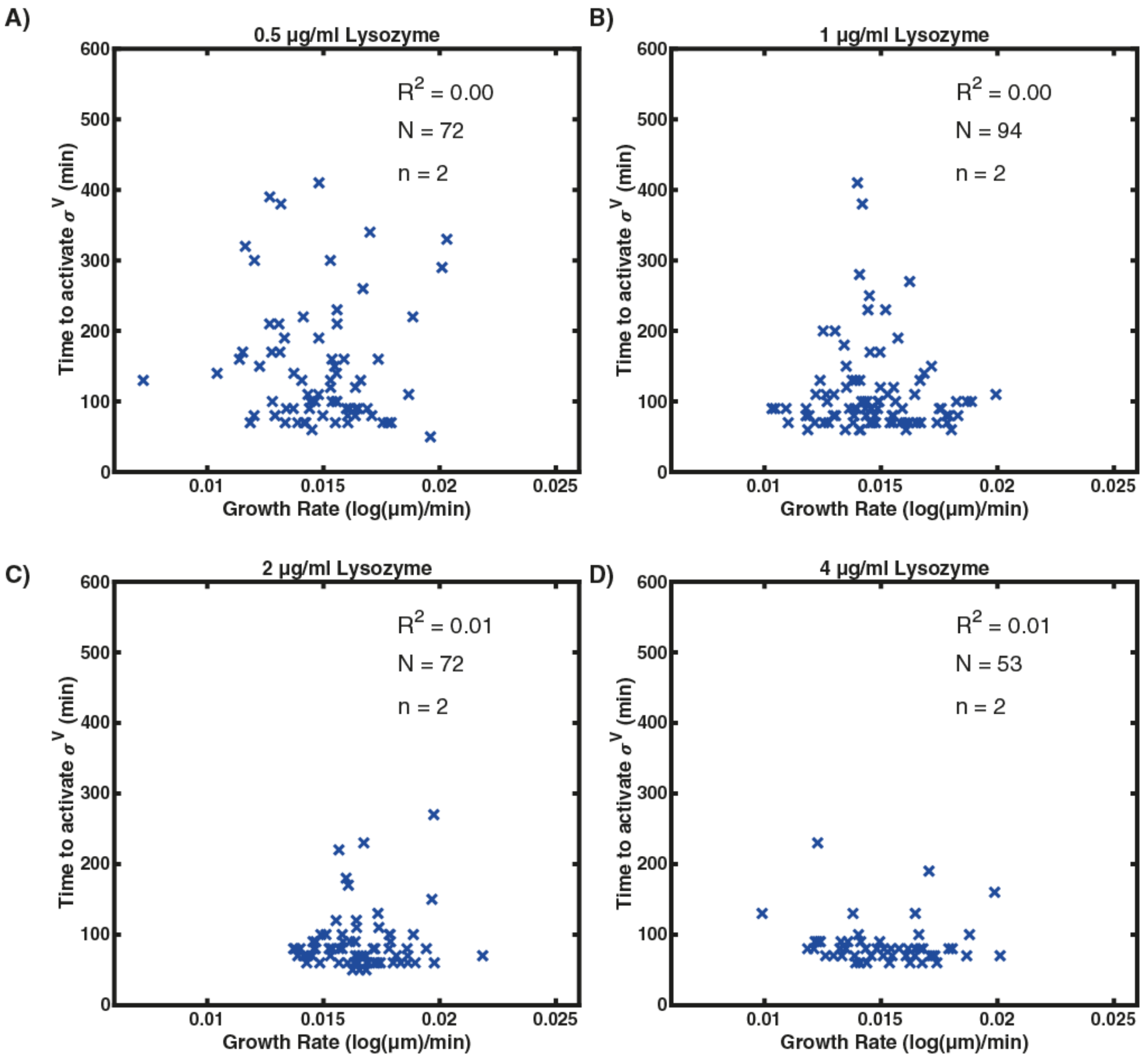

**Figure S11. There is no correlation between growth rate and the activation time.** Figure plots the growth rate at the time of stress addition, against the time to $P_{sigV}$-YFP activation. Each cross corresponds to a single cell. (A) Growth rate vs. activation time for 0.5 µg/ml lysozyme ($R^2$=0.00). (B) Growth rate vs. activation time for 1 µg/ml lysozyme ($R^2$=0.00). (C) Growth rate vs. activation time for 2 µg/ml lysozyme ($R^2$=0.01). (D) Growth rate vs. activation time for 4 µg/ml lysozyme ($R^2$=0.01). Only growth rates during a cell cycle are plotted and not during a division. For more information see methods. Thus N is smaller than in Figure S12.

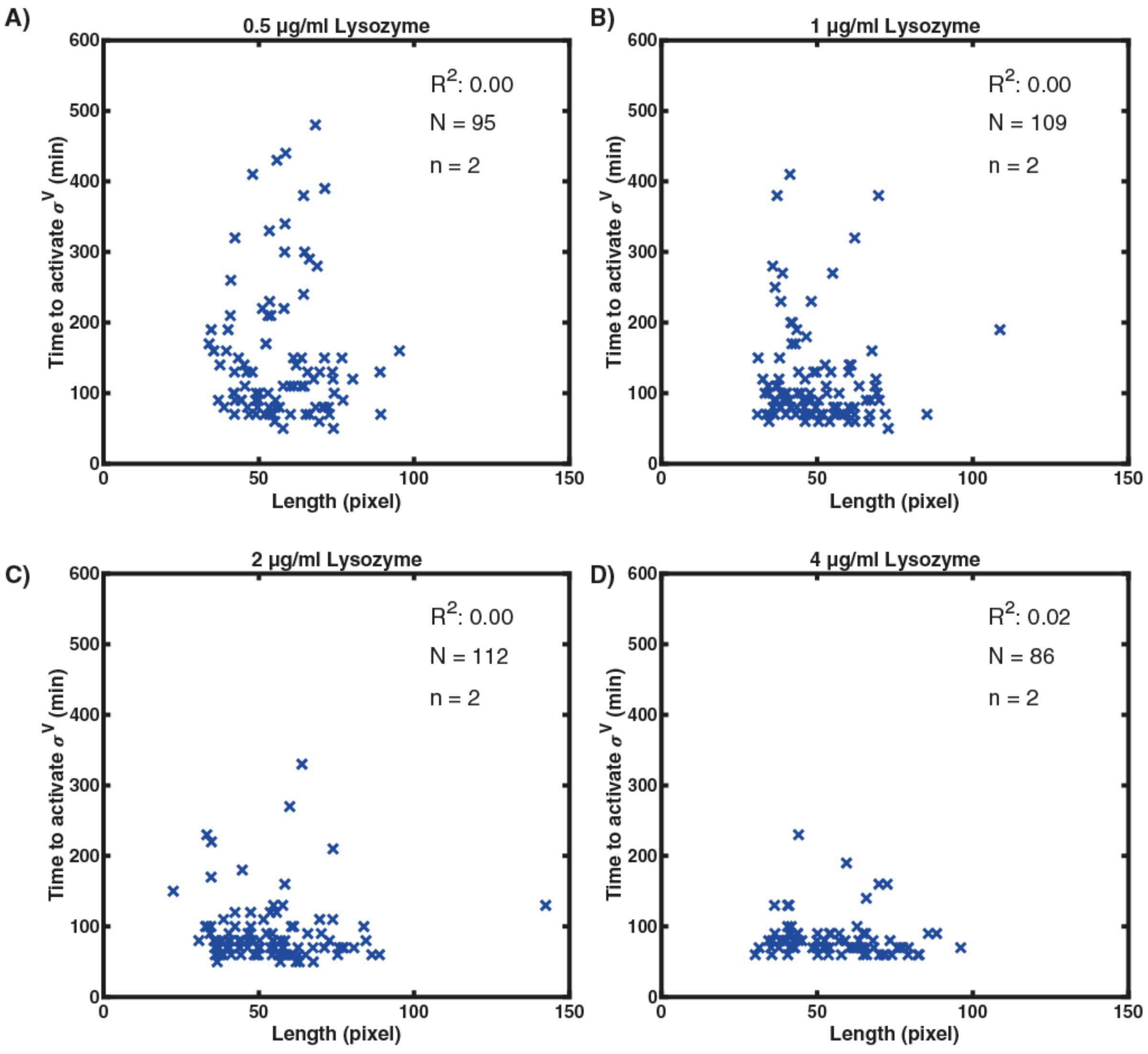

**Figure S12. There is no correlation between cell length and the activation time.** Figure plots the cell length at the time of stress addition, against the time to $P_{sigV}$-YFP activation. Each cross corresponds to a single cell. (A) Cell length vs. activation time for 0.5 µg/ml lysozyme ($R^2$=0.00). (B) Cell length vs. activation time for 1 µg/ml lysozyme ($R^2$=0.00). (C) Cell length vs. activation time for 2 µg/ml lysozyme ($R^2$=0.00). (D) Cell length vs. activation time for 4 µg/ml lysozyme ($R^2$=0.01).

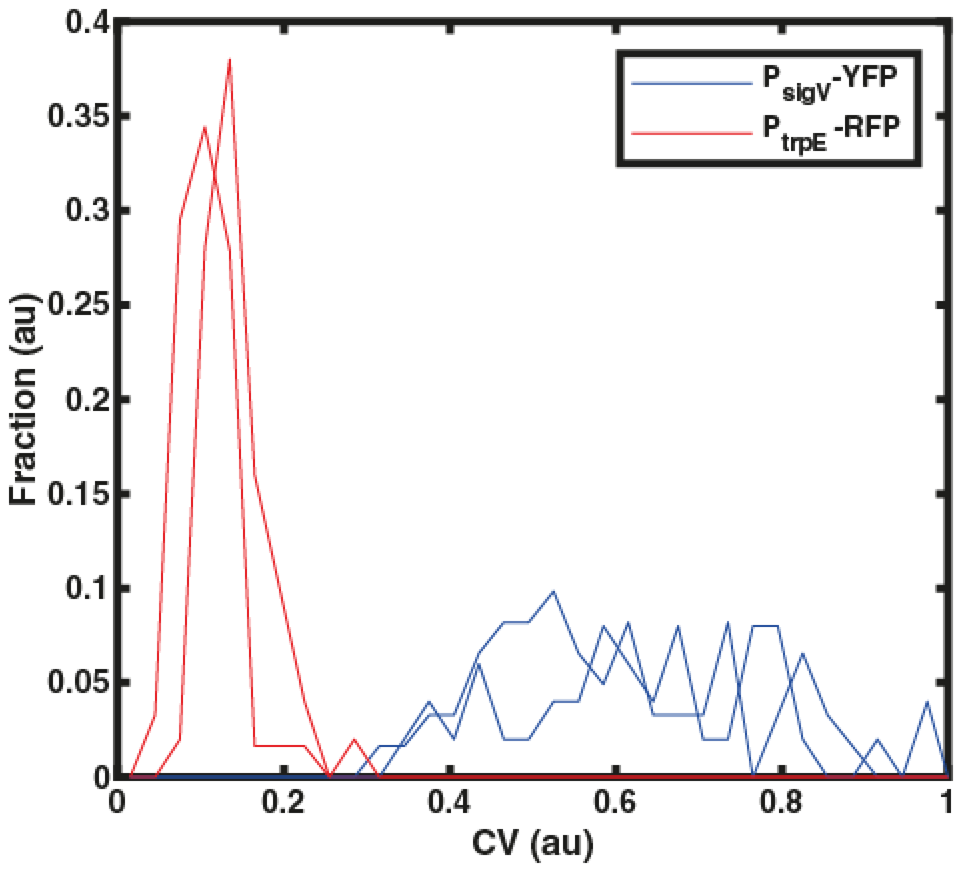

**Figure S13. *sigV* expression displays variability even in absence of stress.** The CV for each mother cell across all timepoints (800 min) was calculated in the absence of lysozyme for $P_{trpE}$-RFP and $P_{sigV}$-YFP . The shown data is from two biological repeats with N ≥ 50 cells.

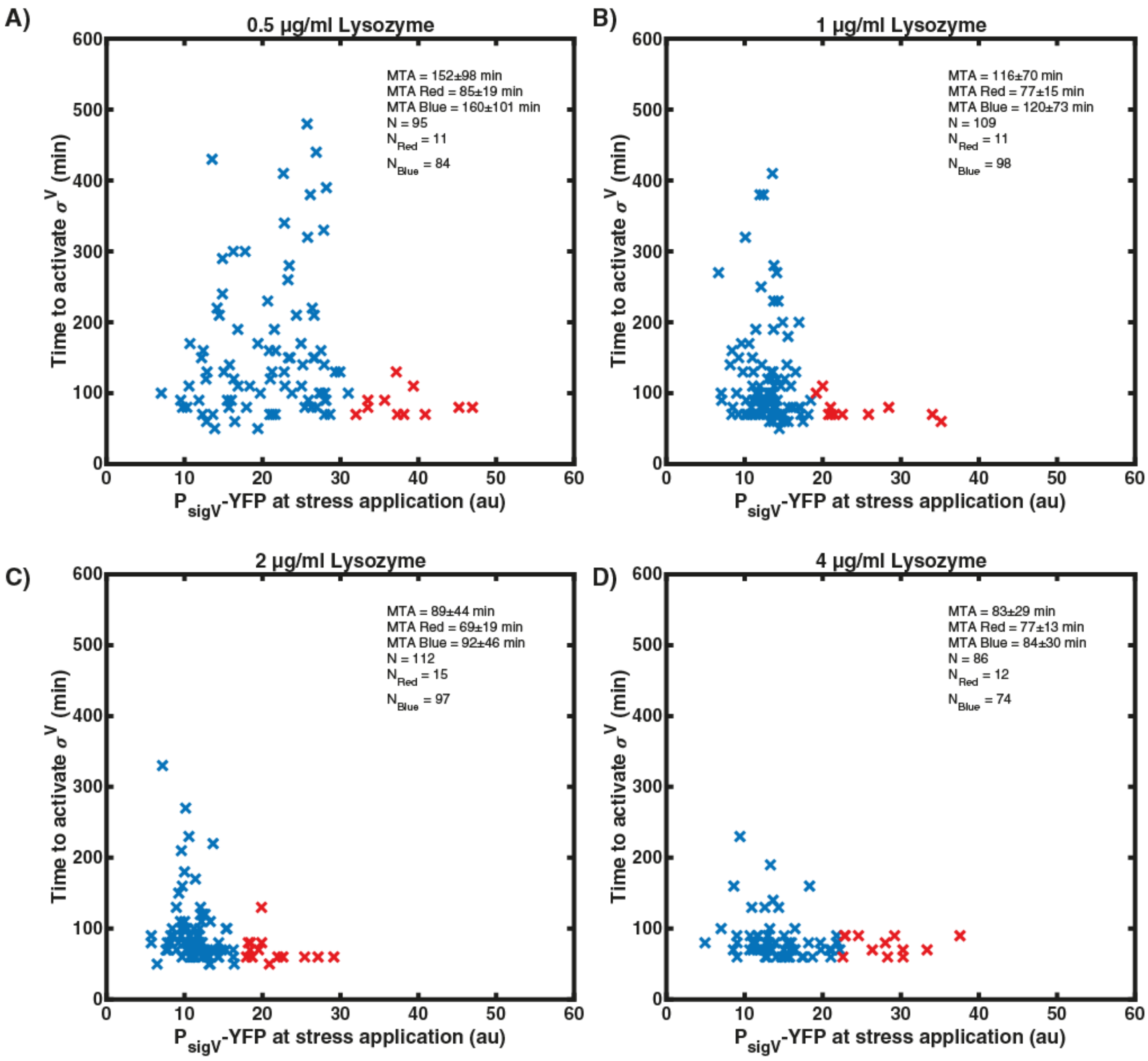

**Figure S14. Cells with high $P_{sigV}$-YFP before stress activate $\sigma^V$ rapidly.** Figure plots $P_{sigV}$-YFP levels at the time of stress addition, against the time to activation. Cells with a YFP fluorescence larger than the mean + SD before stress are marked with red crosses. All other cells are marked with blue crosses. (A) Activation time for 0.5 μg/ml lysozyme. (B) Activation time for 1 μg/ml lysozyme. (C) Activation time for 2 μg/ml lysozyme. (D) Activation time for 4 μg/ml lysozyme. All shown data in the plots is from two biological repeats.

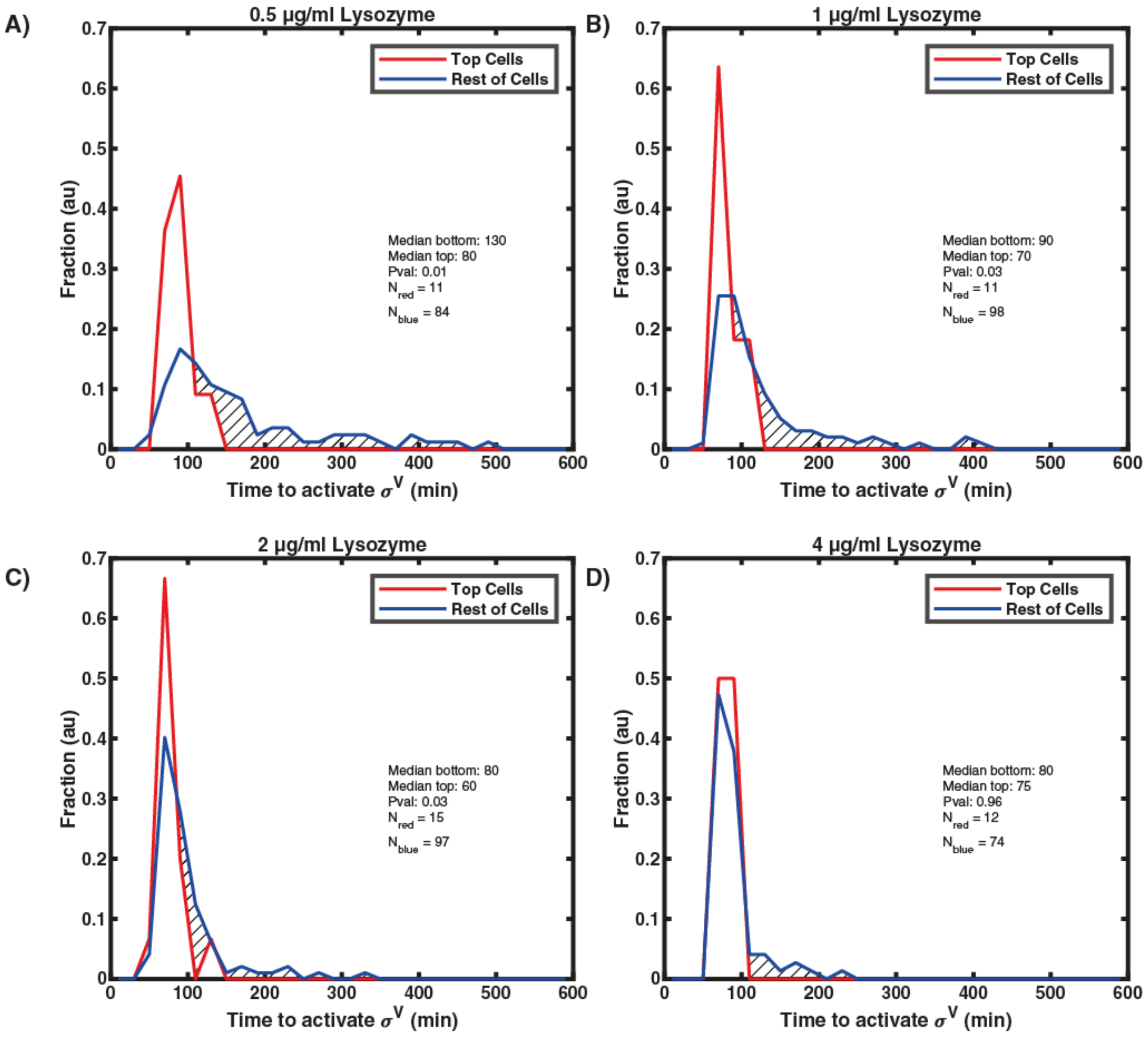

**Figure S15. Cells with higher $P_{sigV}$-YFP levels before stress are more likely to activate $\sigma^V$ instantaneously on lysozyme application.** Cells with a YFP fluorescence larger than the mean + SD are defined as Top Cells (red distribution). All other cells are captured in the blue distribution. The null hypothesis that cells with high $P_{sigV}$-YFP would have the same activation time distribution as cells with low $P_{sigV}$-YFP was rejected for 0.5 μg/ml, 1 μg/ml and 2 μg/ml lysozyme with a p-value of 0.05 using a Kolmogorov–Smirnov test. For more information see Methods. (A) Activation time distributions for 0.5 μg/ml lysozyme. (B) Activation time distributions for 1 μg/ml lysozyme. (C) Activation time distributions for 2 μg/ml lysozyme. (D) Activation time distributions for 4 μg/ml lysozyme. All shown data in the plots is from two biological repeats.

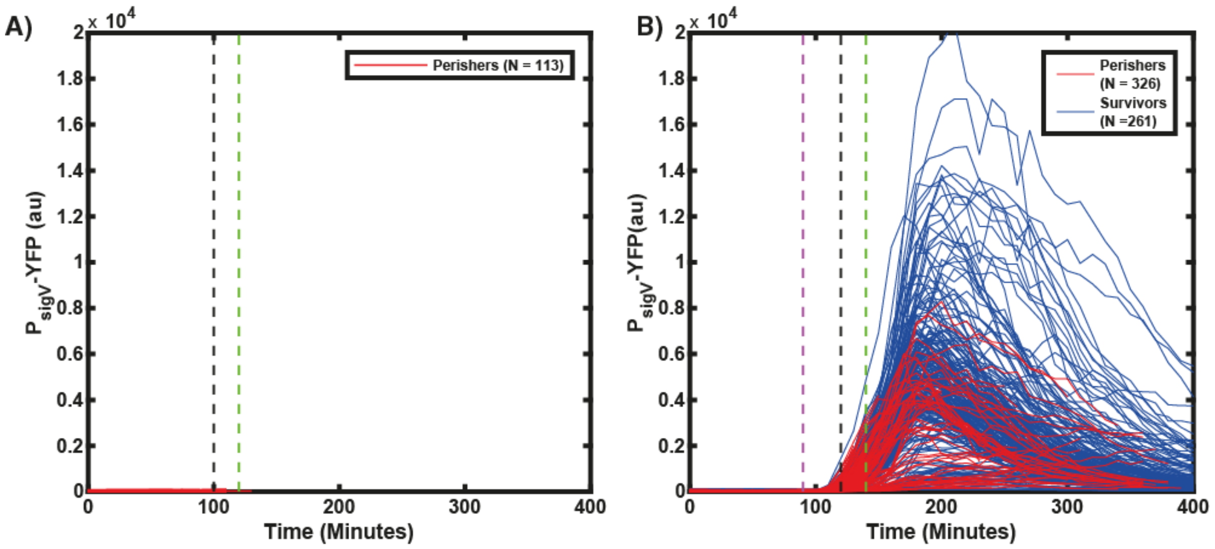

**Figure S16. A sub-lethal stress improves survival to a subsequent lethal stress.** Cells were grown in the mother machine. Each line corresponds to the single-cell time trace of one mother cell. Perishers are defined as cells which die within 280 min (the end of the movie) after the addition of 20 µg/ml lysozyme while survivors continue to grow. (A) All cells die when exposed directly to high lysozyme concentrations. Cells were first grown in SMM and then at the black dashed line exposed to 20 µg/ml lysozyme. After 20 min the lysozyme stress was removed and the media was switched back to SMM (green dashed line). (B) Some cells survive the pulse of 20 µg/ml lysozyme having been exposed to 1 µg/ml lysozyme first. Cells were first grown in SMM before switching to 1 µg/ml lysozyme (magenta dashed line) for 30 min. This short exposure allows cells to activate their $\sigma^V$ pathway and prepare the cell for the subsequent higher stresses. At the black dashed line the media was switched to 20 µg/ml lysozyme for 20 min before switching back to SMM (green dashed line).

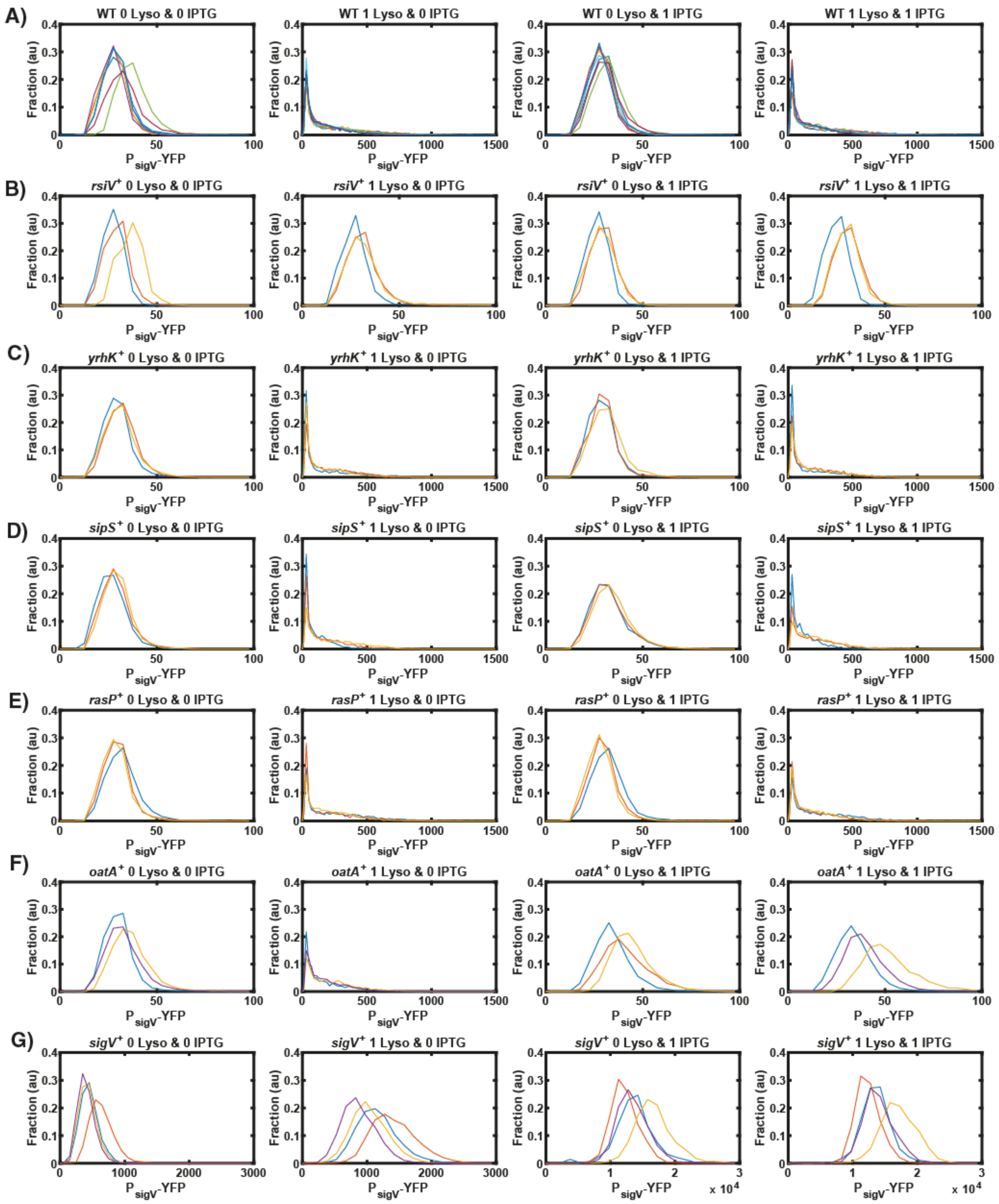

**Figure S17. Overexpression of *yrhK, sipS,* and *rasP* do not change the activation behaviour.** Each column corresponds to a different condition (0 mM IPTG and 0 µg/ml lysozyme, 0 mM IPTG and 1 µg/ml lysozyme, 1 mM IPTG and 0 µg/ml lysozyme, 1 mM IPTG and 1 µg/ml lysozyme). Each row has a different mutant: (A) WT (JLB130), (B) *rsiV*$^+$

(JLB193), (C) *yrhK*$^{+}$ (JLB215), (D) *sipS*$^{+}$ (JLB216), (E) r*asP*$^{+}$ (JLB217), (F) *oatA*$^{+}$ (JLB218), (G) *sigV*$^{+}$ (JLB210). Each coloured histogram corresponds to a different biological repeat (n ≥ 3) and each histogram consists of N >900 cells. For more information on the number of repeats and number of cells please see the supplementary information.

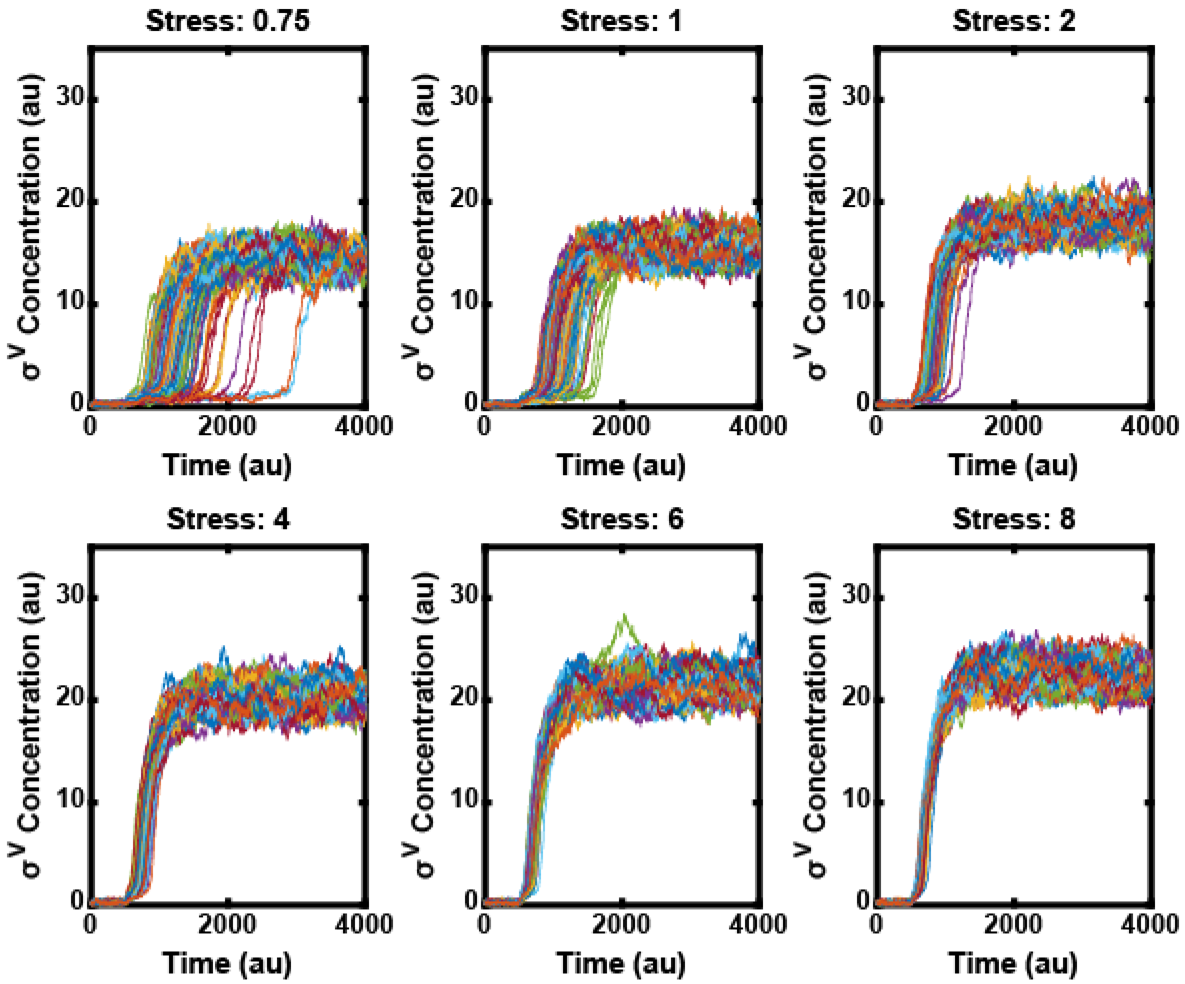

**Figure S18. With increasing stress levels the heterogeneity disappears in the mathematical model.** Each line corresponds to one simulated cell. The stress was added at time point 500 (N=100).

**Figure S18**

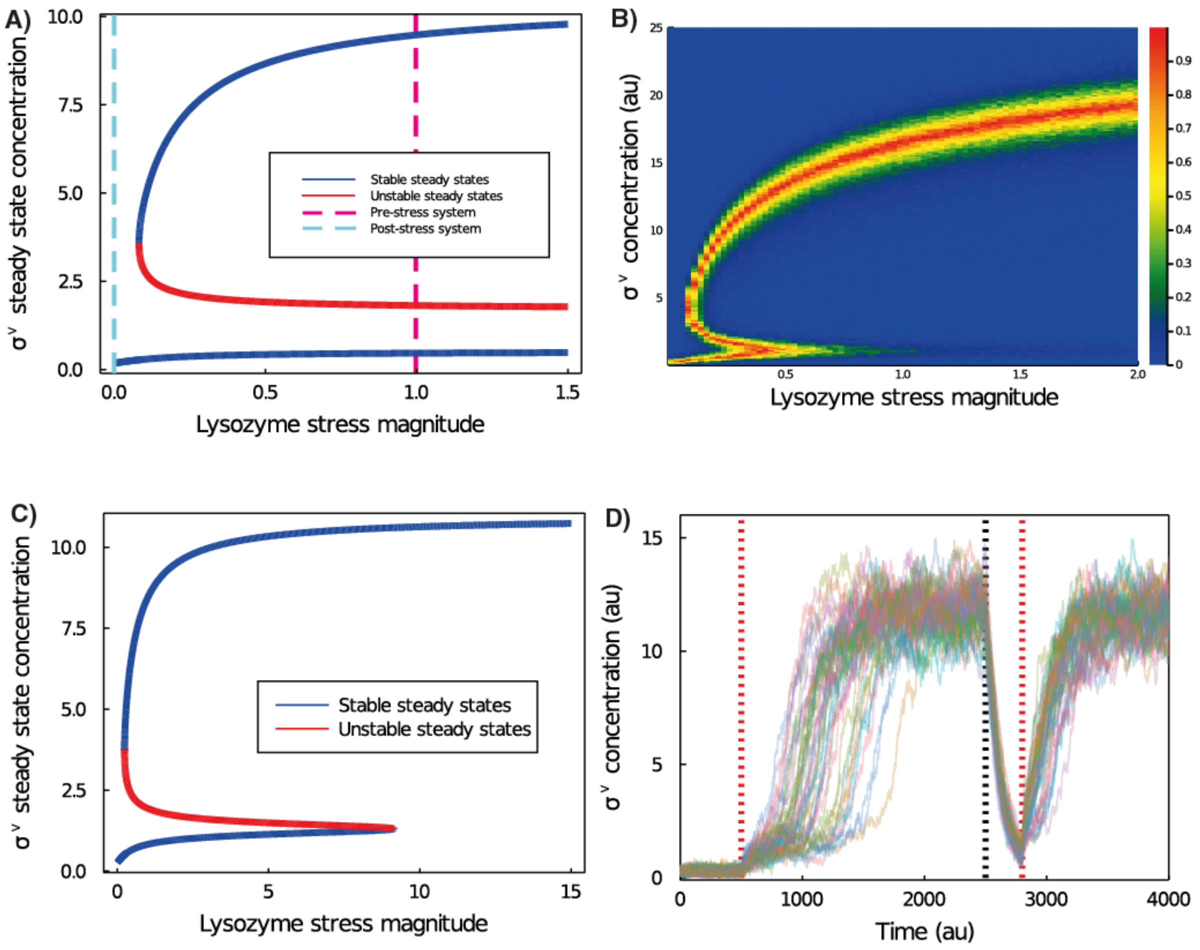

**Figure S19. Deterministic and stochastic bifurcation diagrams showing how the steady states of the model depend on stress input.** (A) Deterministic bifurcation diagram. Without the presence of stress (cyan dashed line) the system is monostable, with only the inactive state present. As stress is added (magenta dashed line), the active state appears. (B) Heatmap shows an approximation of the bifurcation diagram, but for the stochastic implementation of the model. It shows a measure of the stability of the system for each combination of $\sigma^V$ concentration and lysozyme stress. To calculate this, for each lysozyme stress level, initial conditions of $\sigma^V$ concentration were set from 0 and 25 (for 250 values), and then the model was run 100 times for 1000 timepoints for each starting condition. Heat map colour represents the similarity between the probability that the simulation attains a value higher than the initial condition, and the probability that the simulation attains a value lower than the initial condition (the value is 1 when the probabilities are identical, and 0 when either alternative is certain). As stress is increased, the stability of the inactive state (with low $\sigma^V$ concentration) is reduced. This shows how activation times are anti-correlated with stress levels. (C) Alternative deterministic bifurcation diagram. The deterministic version of the model (A) is unable to activate (since the inactive state remains as stress levels approaches infinity). For this slightly different choice of parameters ($v_0$=0.01, K = 3.0, $k_C$=0.35), however, the deterministic model exhibits only a finite region of bistability, allowing for deterministic activation. (D) Simulations from a stochastic model with these modified parameters still exhibit heterogeneous activation, with a memory of stress.

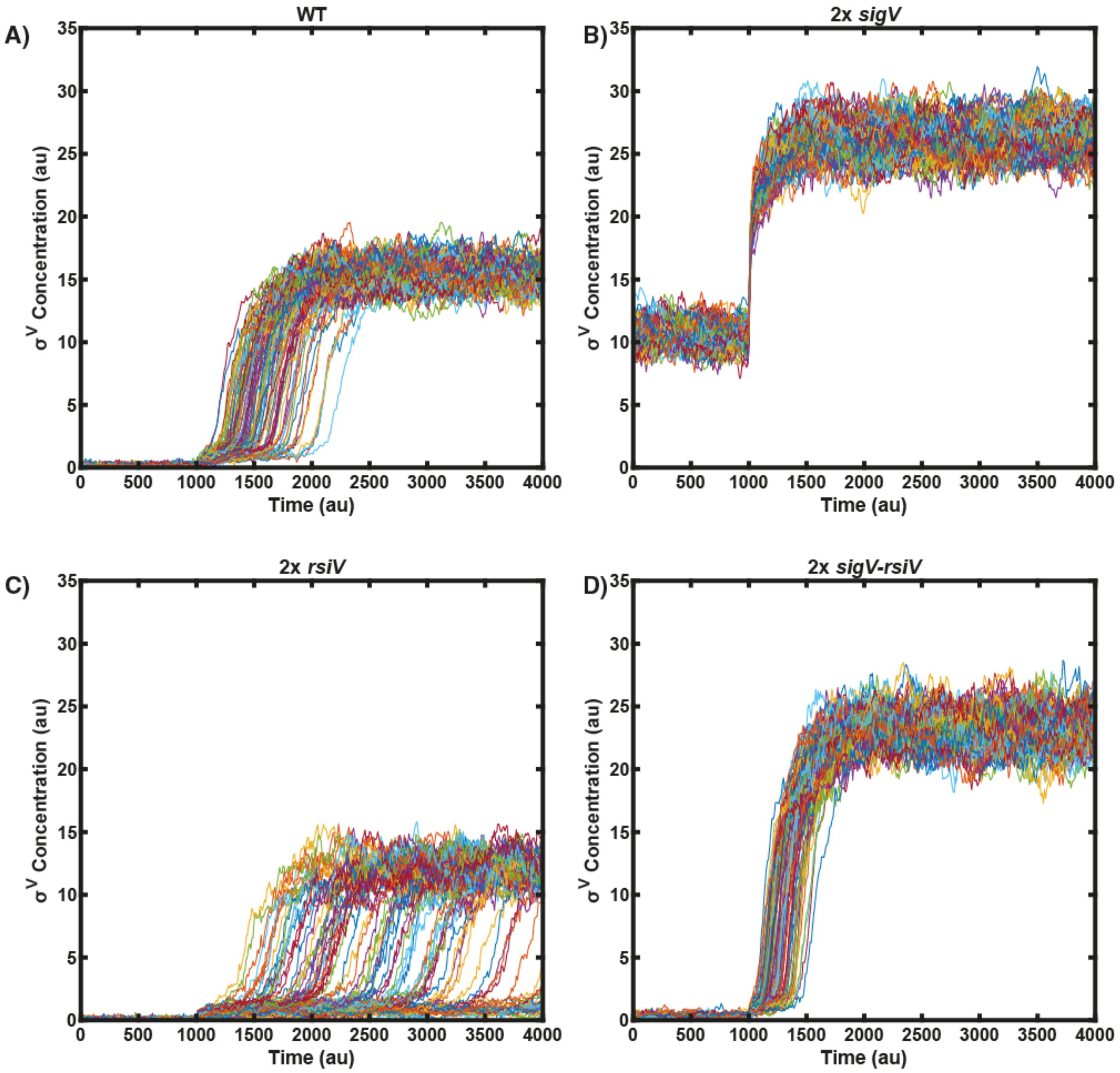

**Figure S20. The model predicts a genetic tunability of the heterogeneity in σ$^V$ response to lysozyme.** In these model simulations a second copy of either *sigv*, *rsiV* or both *sigV* and *RsiV* was introduced (N=100). Stress is added at time 1000. (A) WT, (B) 2x *sigV,* (C) 2x *rsiV,* (D) 2x *sigV-rsiV*

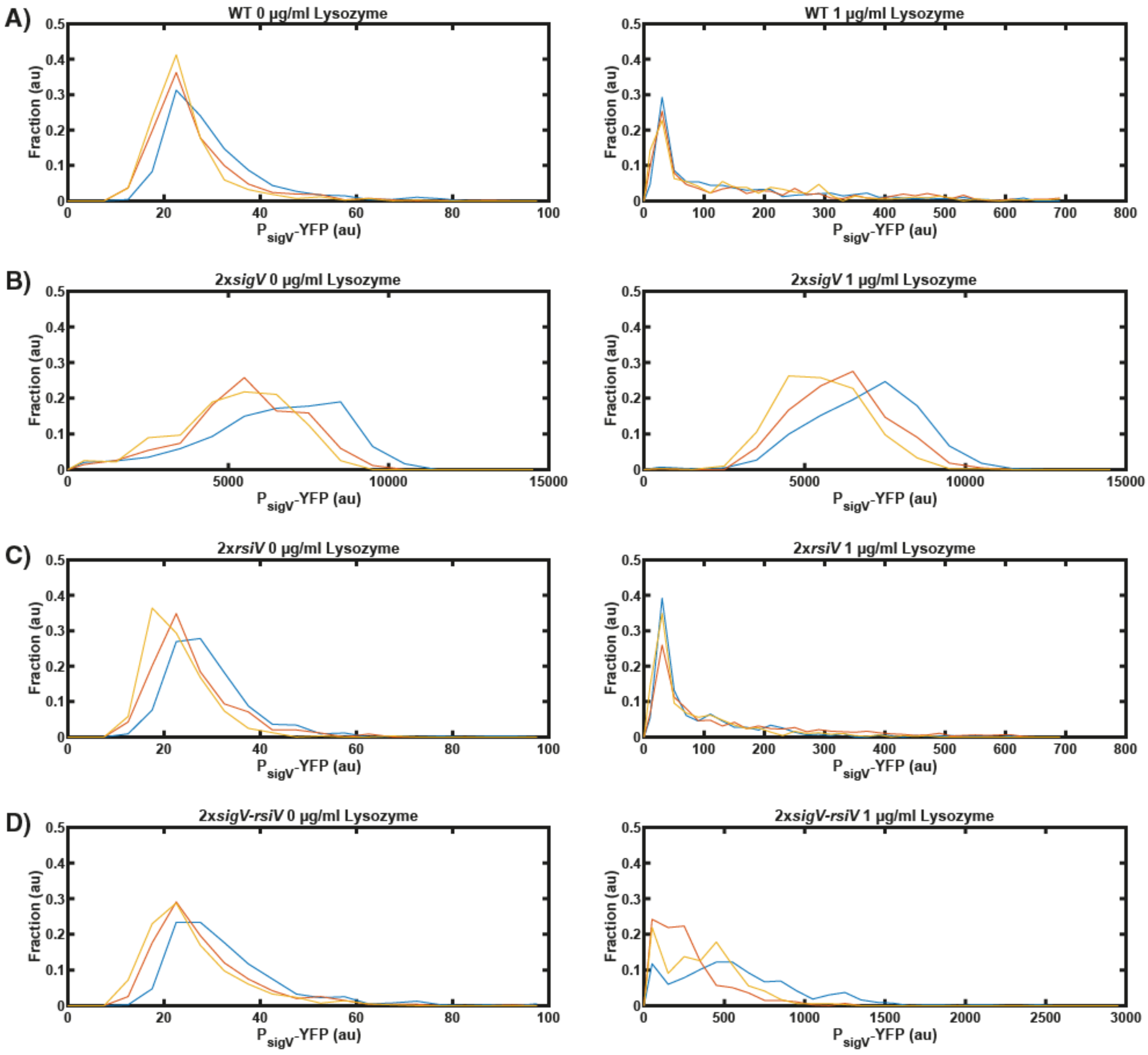

**Figure S21. Snapshots confirm that the $\sigma^V$ response to lysozyme is genetically tunable.** The first column corresponds to 0 µg/ml lysozyme and the second column corresponds to 1 µg/ml lysozyme. Each row is a different mutant: (A) WT (JLB130), (B) 2x *sigV* (JLB210), (C) 2x *rsiV* (JLB212), (D) 2x *sigV-rsiV (JLB211)*.

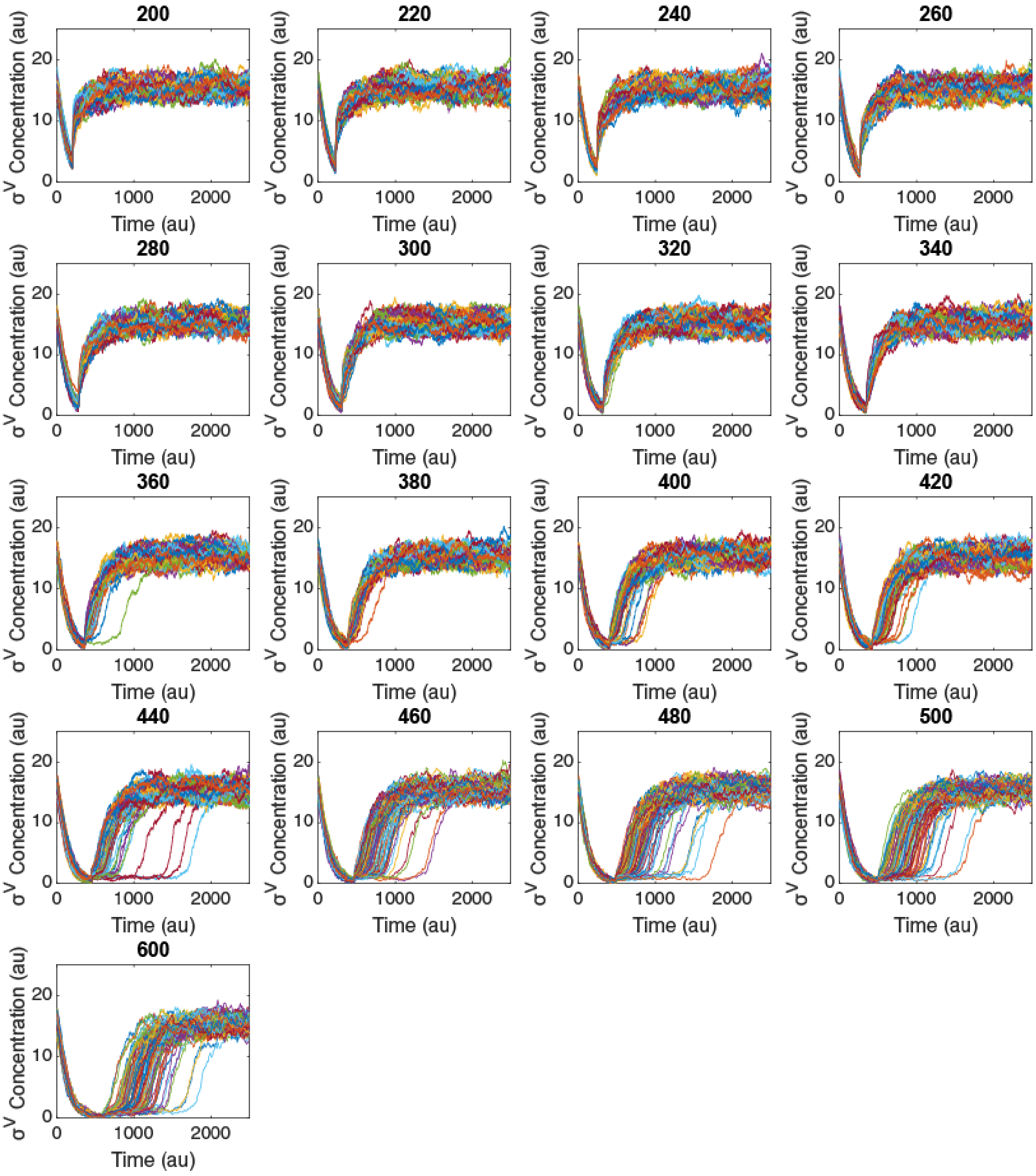

**Figure S22. The memory effect decreases with increasing stress holidays in the model.** Each line corresponds to one simulated cell. At time 0 the stress was removed and reapplied at time points 200 - 600 (N=100 for each panel).

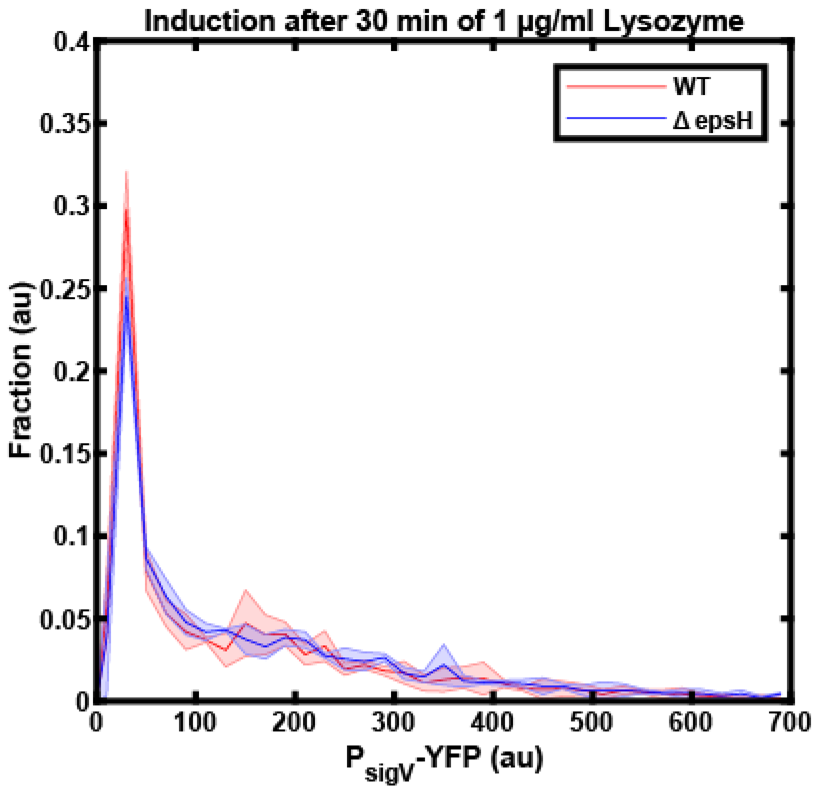

**Figure S23. WT and ΔepsH have the same $P_{sigV}$-YFP activation in snapshots.** WT (JLB130) and ΔepsH (JLB221) were grown in SMM with 1 μg/ml lysozyme for 30 min and then investigated with snapshots (n=3).

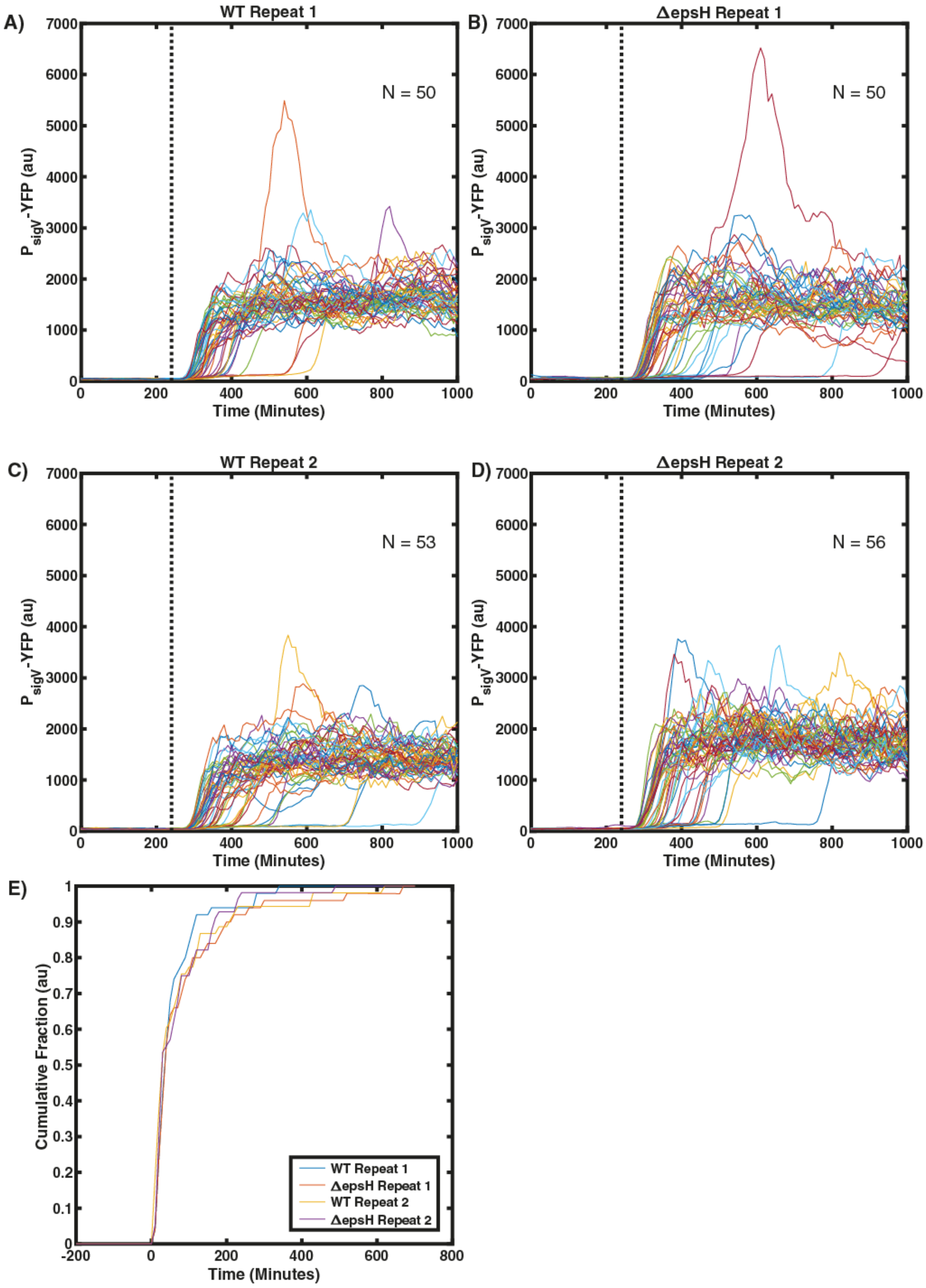

**Figure S24. WT and ΔepsH have similar $P_{sigV}$-YFP activation in movie experiments.**

(A-D) (JLB130) and ΔepsH (JLB221) were grown in SMM in the mother machine microfluidics device for 240 h before switching the media to SMM with 1 μg/ml lysozyme. (E) Each line corresponds to the fraction of cells with a $P_{sigV}$-YFP larger than its half maximum at a given time point. WT (JLB130) and ΔepsH (JLB221) were grown in the mother machine in SMM for 240 min before switching to 1 μg/ml lysozyme (n=2). For more information on the repeats and number of cells please the the supplementary text.

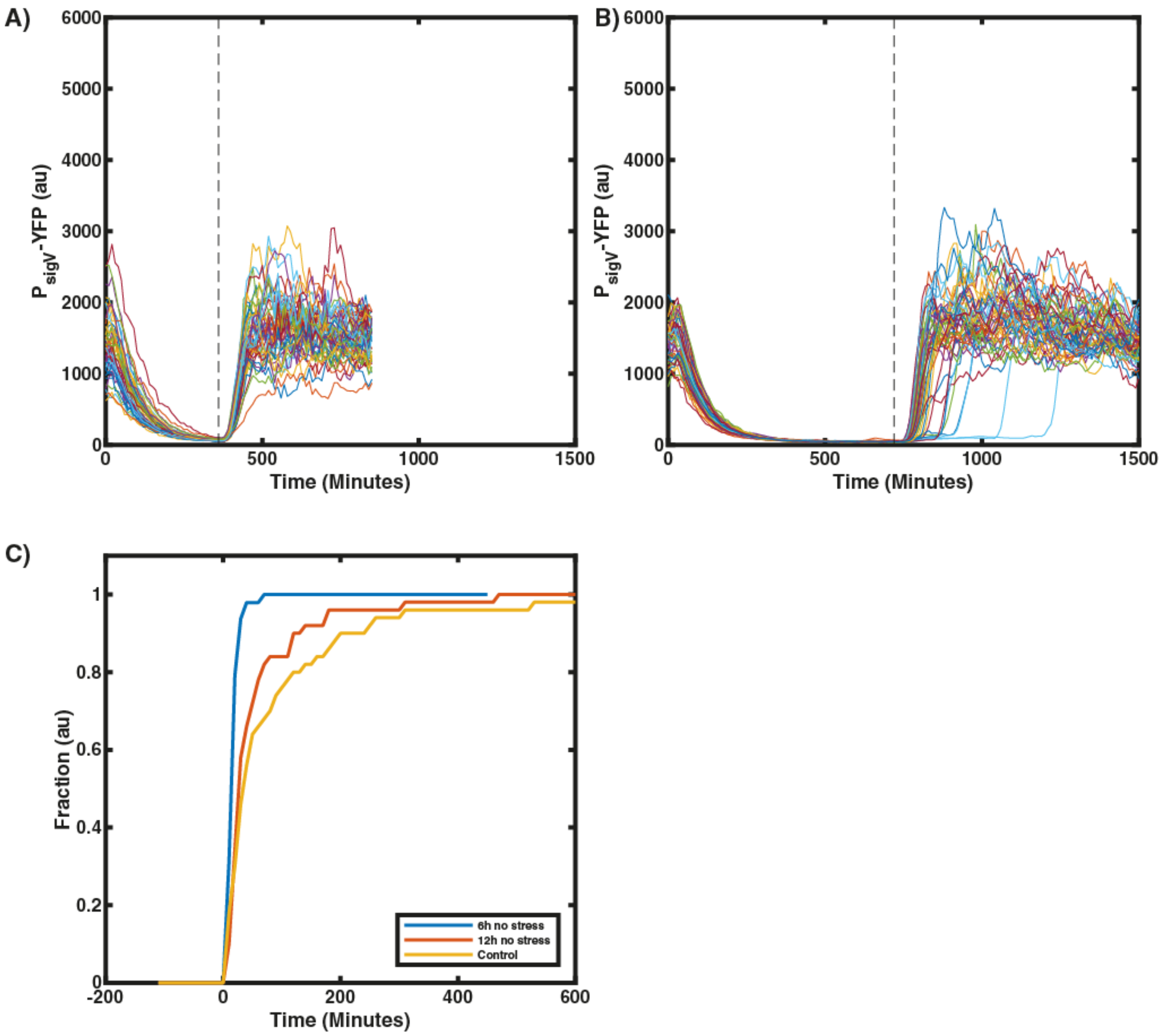

**Figure S25. Repeat experiment also shows a memory in $\sigma^V$ after a stress break.** The stress was removed at time 0 min and reapplied after 6h or 12h. Each line corresponds to one mother cell. (A) 6h stress break. (B) 12h stress break. (C) This figure shows the fraction of cells with PsigV-YFP larger than their half maximum.

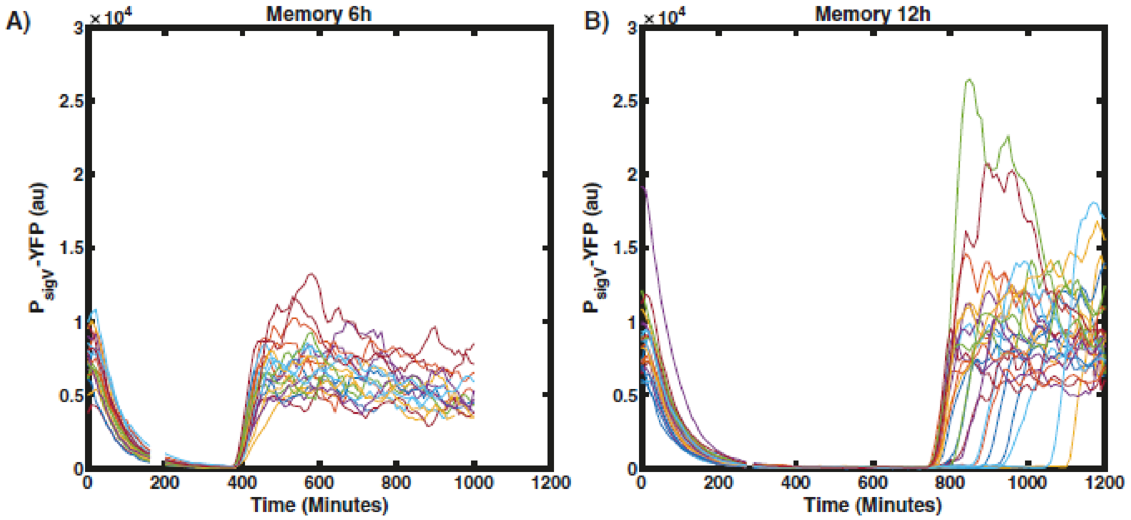

**Figure S26. The memory dynamics are independent of the design of the mother machine microfluidic device.**

JLB130 was grown in the mother machine microfluidic device with a design from the Paulsson lab (Norman et al. 2013) rather than our standard mother machine design based on the chip from the Jun lab (Wang et al. 2010). (A) After a 6h long break in stress all cells responded homogeneously to the reapplication of 1 µg/ml lysozyme (N=20). (B) Cells grown in the mother machine were exposed to a 12h long stress holiday. After the break in stress the heterogeneity in $\sigma^V$ activation reappeared (N=26). During the gaps in the traces the cells were out of focus. (n=1).
